# MITRE: predicting host status from microbiota time-series data

**DOI:** 10.1101/447250

**Authors:** Elijah Bogart, Richard Creswell, Georg K. Gerber

## Abstract

Longitudinal studies are crucial for discovering casual relationships between the microbiome and human disease. We present Microbiome Interpretable Temporal Rule Engine (MITRE), the first machine learning method specifically designed for predicting host status from microbiome time-series data. Our method maintains interpretability by learning predictive rules over automatically inferred time-periods and phylogenetically related microbes. We validate MITRE’s performance on semi-synthetic data, and five real datasets measuring microbiome composition over time in infant and adult cohorts. Our results demonstrate that MITRE performs on par or outperforms “black box” machine learning approaches, providing a powerful new tool enabling discovery of biologically interpretable relationships between microbiome and human host.

## Introduction

The human microbiome is highly dynamic on multiple timescales, changing dramatically during development of the gut in childhood, with diet, or due to medical interventions [1]. Recently, a number of longitudinal studies have been undertaken, seeking to link changes in the microbiota over time with medical interventions such as delivery by Cesarean section [2], dietary changes [3], or antibiotic treatment [4], or with disease outcomes in the host such as type 1 diabetes [5], dietary allergies [6], premature delivery [7, 8], necrotizing enterocolitis [9, 10] and infection [11, 12].

Deriving maximally useful information from these studies requires computational methods that can simultaneously identify patterns of change in the microbiome and link these patterns to the host’s status (e.g. disease outcome, presence or absence of an intervention, etc.) Moreover, such computational methods must contend with numerous challenges inherent to microbiome time-series data, including measurement noise, sparse and irregular temporal sampling, and inter-subject variability.

To overcome the challenges inherent in analyzing microbiome time-series data to predict host status, we developed MITRE, a computational model that infers human-interpretable predictive rules from high-throughput microbiome time-series data, implemented in an open-source software package (https://github.com/gerberlab/mitre/). MITRE falls into the general category of Bayesian supervised machine learning classifiers: the algorithm uses a training dataset of microbiota time-series and host statuses (supervised learning) to learn a probability distribution (Bayesian inference) over a set of alternative models that predict the status of a host from associated microbiome data (classification). Bayesian approaches are powerful, because they provide principled estimates of uncertainty throughout the model, which is an especially important feature in biomedical applications where noisy inputs are the norm. We note that another rule-based method, association rule mining (ARM), has recently been applied to analyzing microbiome data in a different context (finding interaction patterns among OTUs) [13]. Although ARM has some commonalities with Bayesian rule learning approaches, ARM methods tend to employ user-based cut-offs and heuristics, rather than principled probabilistic methods, as their primary function is to mine large databases for putative interactions, rather than build predictive models. Further, unlike Bayesian models, ARM methods do not incorporate prior knowledge, as their focus is mining from large databases.

In previous work, we presented the MDSINE [14] algorithm, which infers dynamical systems models from microbiome time-series data in order to predict population dynamics of the microbiome over time. Our present work, MITRE, addresses a different question: can we predict the status of the host *given* microbiome time-series data. From the machine learning perspective, MDSINE is an unsupervised model, whereas MITRE is a supervised model. The key distinction is that MDSINE models the microbiome, whereas MITRE does not, and instead models host outcomes. Like other supervised models, MITRE focuses on finding only the essential features (in this case, microbial clades and relevant time-windows) to explain the outcome, rather than attempting to explain the microbiome data itself. This architecture is ideal for highly heterogeneous datasets with many “distractors,” which are the reality for longitudinal studies of the human microbiome. Supervised machine learning classifiers are employed in many biomedical predictive modeling applications, including forecasting (predicting a future outcome, such as onset of disease, based on past data) and diagnosis or subtyping (predicting which category a subject belongs to based on all available data).

MITRE’s unique contributions are its modeling of the special properties of microbiome time-series data (phylogenetic and temporal relationships), and its emphasis on producing human-interpretable predictors. This latter capability is in contrast to various generic “black box” machine learning methods that have been applied to analyzing static microbiome data, such as random forests [6, 15–17], which may achieve high predictive accuracy but do not yield easily human-interpretable models; interpretability is especially challenging for such models in the context of time-series analyses, given repeated measurements and the fact that relevant dynamics may occur at multiple time-scales. In the following sections, we introduce the MITRE framework, then provide benchmarking results of MITRE versus comparator methods on semi-synthetic data and five real microbiome time-series datasets, and finally illustrate through examples MITRE’s exploratory data analysis capabilities and how these can help extract biological insights.

## Results

### Conceptual overview of the MITRE model and software

Figure 1 provides an overview of the MITRE framework. MITRE takes as input: (1) tables of microbial abundances, typically operational taxonomic units (OTUs) from 16S rRNA amplicon sequencing or species mappings from metagenomic data, measured over time for each host, (2) the status of each host (e.g., phenotype A or phenotype B), (3) an optional set of static covariates for each host (e.g., gender), and (4) placements of the microbes on a reference phylogenic tree [18]. MITRE then automatically learns from the data predictive models that can be expressed as a set of conditional statements, or human-readable rules, about time-localized patterns of change in the abundances of groups of phylogenetically related organisms. Weighted sums of the truth values of the rules are used to predict the status of each host. The MITRE software package also provides a graphical user interface (GUI) for interactive visualization of the output, which summarizes the predictive models learned from the data.

**Figure 1:**
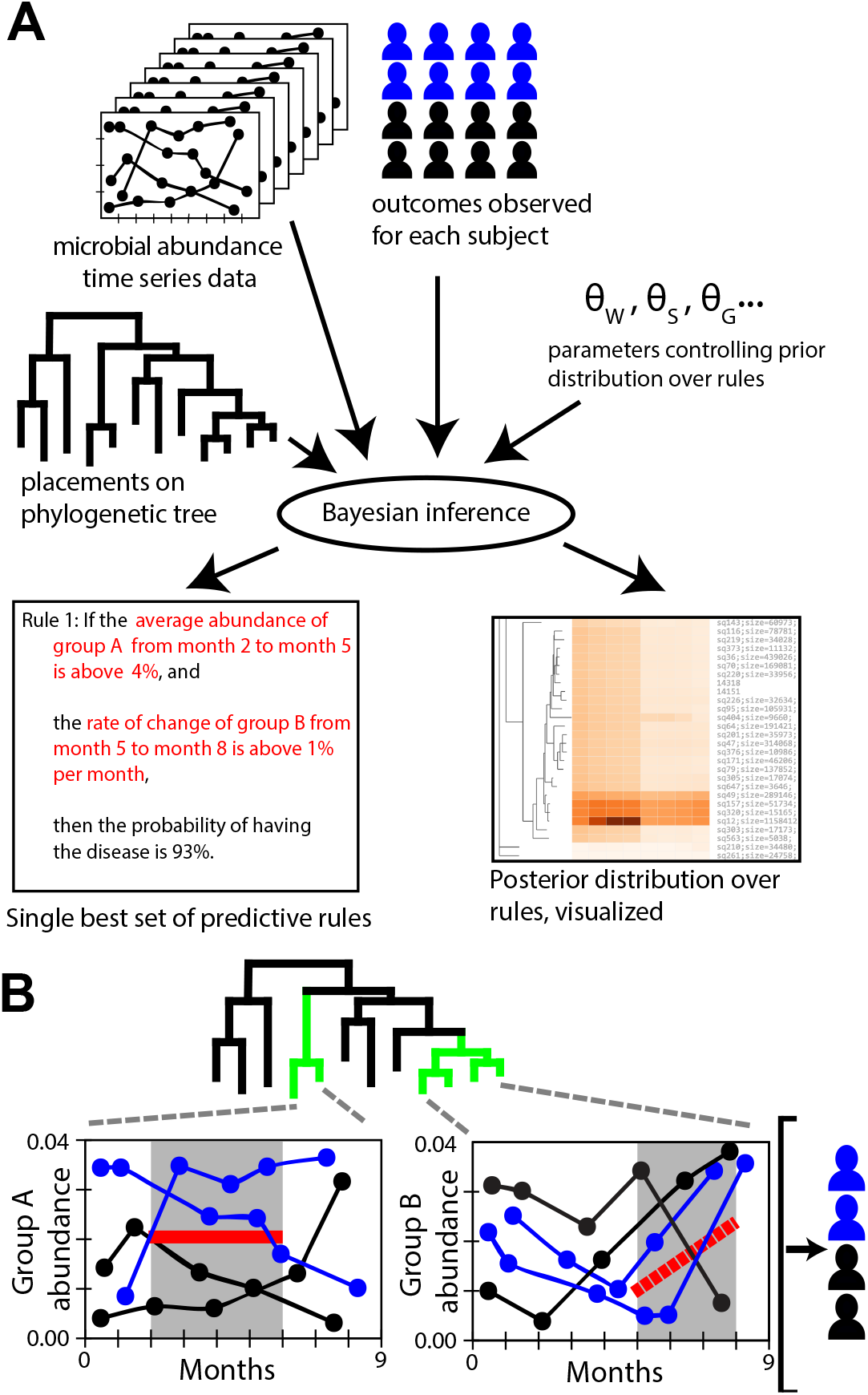
MITRE learns human-interpretable rule-based models predicting host status from microbiota time-series data. Rules operate on automatically learned time-periods and groups of phylogenetically related microbes. **(A)** Schematic of the MITRE analysis pipeline, resulting in a single best predictive model as well as a distribution over alternative models that can be interactively explored. **(B)** Schematic of example rule in **(A)** applied to hypothetical data. Here, two subjects satisfy both the condition on the average abundance of microbe group A and the rate of change of abundance of group B.

To be more precise, a MITRE model consists of a baseline probability of a default host status plus a set of zero or more rules. Each rule is a conjunction of one or more *detectors*—conditional statements about bacterial abundances in the form “between times *t_0_* and *t_1_*, the average abundance of bacterial group *j* is [above/below] threshold *θ_l_”* or “between times *t_0_* and *t_1_*, the slope of the abundance of bacterial group *j* is [above/below] threshold *θ_l_*”—together with a multiplicative effect on the odds of the outcome of interest if all the detectors are satisfied.

As a simple example, a MITRE model predicting the odds of an infant developing a disease in the first year of life might be:

- If, from month 2 to month 5, the average relative abundance of bacterial clade A is above 4.0%, and from month 5 to month 8, the relative abundance of bacterial clade B increases by at least 1.0% per month, the odds of disease increase by a factor of 10;
- If, from month 3 to month 10, the average relative abundance of OTU C is less than 9.5%, the odds of disease decrease by a factor of 2;
- The baseline probability of disease is 22.0%.

Figure 1B schematically illustrates the application of a rule set to hypothetical data. To predict the probability that an individual will develop the disease, the effects of each rule satisfied by that individual’s microbiome data are combined with the baseline disease probability.

A comprehensive pool of possible detectors is generated automatically at the beginning of a MITRE analysis, including detectors that apply to average values and rates of change of clades at all levels on the phylogenetic tree of observed bacteria at as many time windows as the temporal resolution of the data will allow (see Methods.) By combining detectors from this pool, rules in a MITRE model can capture rich temporal patterns, but still remain human-interpretable because each component rule is easy to understand.

MITRE is a nonlinear model, which has a number of advantages over a linear model. Most obviously, nonlinear models can capture effects such as thresholds/saturation or interactions between variables. A more subtle issue, particularly relevant to microbiome ratio-based data, is that linear models introduce mathematical/statistical difficulties when analyzing compositional or proportional data. While special constraints are needed for linear models to overcome these difficulties [19], nonlinear models do not suffer from the same limitations because they can inherently learn nonlinear transformations of the data. It is particularly straightforward to understand the transformations produced by the MITRE detectors described above: the learned thresholds effectively discretize ratio data into distinct levels, a transformation that renders the data mathematically non-compositional.

The MITRE framework is fully Bayesian, meaning it learns a distribution over models, called the posterior probability distribution, which takes into account both the input data as well as prior information provided. In the case of MITRE, this prior information favors by default parsimonious explanations, i.e., short sets of simple rules. Importantly, the default prior is designed to favor the empty rule set, or the case in which the baseline odds only are used to predict host status, and the microbiome data plays no role. This feature of the default prior is designed to guard against over-fitting. Moreover, through the formalism of Bayes factors (see Methods), MITRE provides a quantitative measure for the evidence favoring no association between the microbiome data and host status or the alternative of any rule sets that predict such a relationship; this feature allows the user to rigorously evaluate whether sufficient signal is present in the microbiome data to predict host status. Of note, this measure in effect incorporates multiple “hypotheses” simultaneously (as the inference procedure explores the entire space of possible rule sets at once.) Additional Bayes factors allow the user to assess the evidence that each particular bacterial clade or OTU is associated with the host status, by comparing the evidence for a model in which no detector in the rule set applies to the clade of interest to a model in which at least one detector does apply to the clade.

The MITRE software approximately infers the posterior probability distribution using a custom Markov Chain Monte Carlo algorithm, and reports a *point estimate* of the single best rule set as well as a summary of the distribution, which the user may investigate interactively with the provided GUI. To make predictions for new data, either the point estimate or an *ensemble* of multiple rule sets, weighted according to their posterior probabilities, may be used.

### Benchmarking against standard machine learning methods: semi-synthetic data

We used semi-synthetic data to compare the cross-validated predictive performance of MITRE to two popular standard machine learning methods, random forests and L1-regularized logistic regression, which have been widely used to analyze data from static microbiome studies. We tested the performance of MITRE and the comparator methods by training each method on a subset of the available data and then classifying the remaining data according to the model learned. For each method, performance was evaluated using the F1-score (harmonic mean of precision and recall) under crossvalidation, converting modeled probabilities of outcomes to binary predictions by applying a threshold at probability 0.5.

We simulated data from a real dataset using a parametric bootstrapping-type procedure, in which models of microbiome dynamics were employed to interpolate the real data and inject temporal perturbations into microbial clades to simulate a “disease” host phenotype (see Methods for details). The real dataset [2] we bootstrapped from tracked the gut microbiome composition from birth to two years of age in a cohort of U.S. infants; we chose this dataset because it was among the densest and most regularly sampled of available time-series datasets, and also studied a relatively large number of subjects. To gain insight into the predictive performance of the different methods, we simulated data with varying numbers of subjects or time-points, and one or two temporal perturbations to microbial clades to simulate subjects with a “disease.” We assumed the perturbations occur over a limited but unknown time-period during the study (∼20% of the study duration), which represents a challenging but biologically relevant scenario for analysis. The ranges of subjects and time-points simulated correspond approximately to those in real studies, and thus provide insight into the performance of the algorithms on realistically sized studies. Note that MITRE is a supervised machine learning method, which does not directly model the covariates (microbiome data.) Thus, our data simulation procedure is unrelated to the underlying MITRE model and not expected to introduce bias in favor of our method.

To provide as reasonable a comparison as possible, we used the average abundance of each OTU in a series of time windows as input to the comparator algorithms. It is important to note that there are no prior methods available that were specifically designed for supervised learning from microbiome time-series data. In fact, the comparator algorithms we implemented themselves represent an advance over the state-of-the-art for many studies. To date, most studies have employed an *ad hoc* strategy of manually identifying time windows of interest within the experiment, and then testing for differential abundance of pre-specified groups of taxa in each time window separately. Such an approach has significant limitations. For example, effects occurring outside defined windows or across their boundaries may not be detected; and, analyzing each time window/taxon pair independently significantly reduces statistical power and precludes the discovery of interactions across taxa or sequences of events across multiple time windows.

Our results on semi-synthetic data over a range of scenarios (Figure 2A-2D) demonstrate superior cross-validated predictive performance of MITRE compared to the other methods. Several interesting trends are also evident from these results. First, the MITRE ensemble and point methods have similar performance, except for in the setting of low numbers of subjects, which is likely simply a stochastic effect since all the methods in that setting have poor performance. The similar performance of the MITRE point and ensemble methods is very encouraging from the interpretability perspective, since the point method yields a single, human-interpretable rule set. Second as expected, all of the methods improve in performance with increasing numbers of subjects, with eventual plateauing of gains in performance at a level ultimately limited by noise in the data. Third, there is also improvement in performance with increasing numbers of time-points, but this improvement is less impressive. This phenomenon can be partially explained by our assumption in generating the semi-synthetic data that perturbations corresponding to the “disease” phenotype occur over a limited time-period during the study. Thus, sampling of more time-points outside the perturbation period provides only limited additional information useful for prediction. Fourth and finally, we also see generally worse performance of all methods in the more complex setting of two perturbations in the “disease” cases, particularly with limited numbers of subjects or time-points. Interestingly, Random Forests outperforms L1-regularized logistic regression in the two-perturbation case in the setting of low numbers of subjects or time-points, while the opposite is true in the one-perturbation case, which may be due to Random Forest’s capacity to handle non-linearities. In any event, MITRE, which models non-linearities through conjunctions in rules, consistently outperforms the other two methods in this setting as well. Overall, our results demonstrate that MITRE, a method specifically tailored for analyzing microbiome time-series data, outperforms generic machine learning methods. Moreover, we provide a simulation and testing platform for users to investigate questions relevant to particular microbiome time-series datasets in the future.

**Figure 2:**
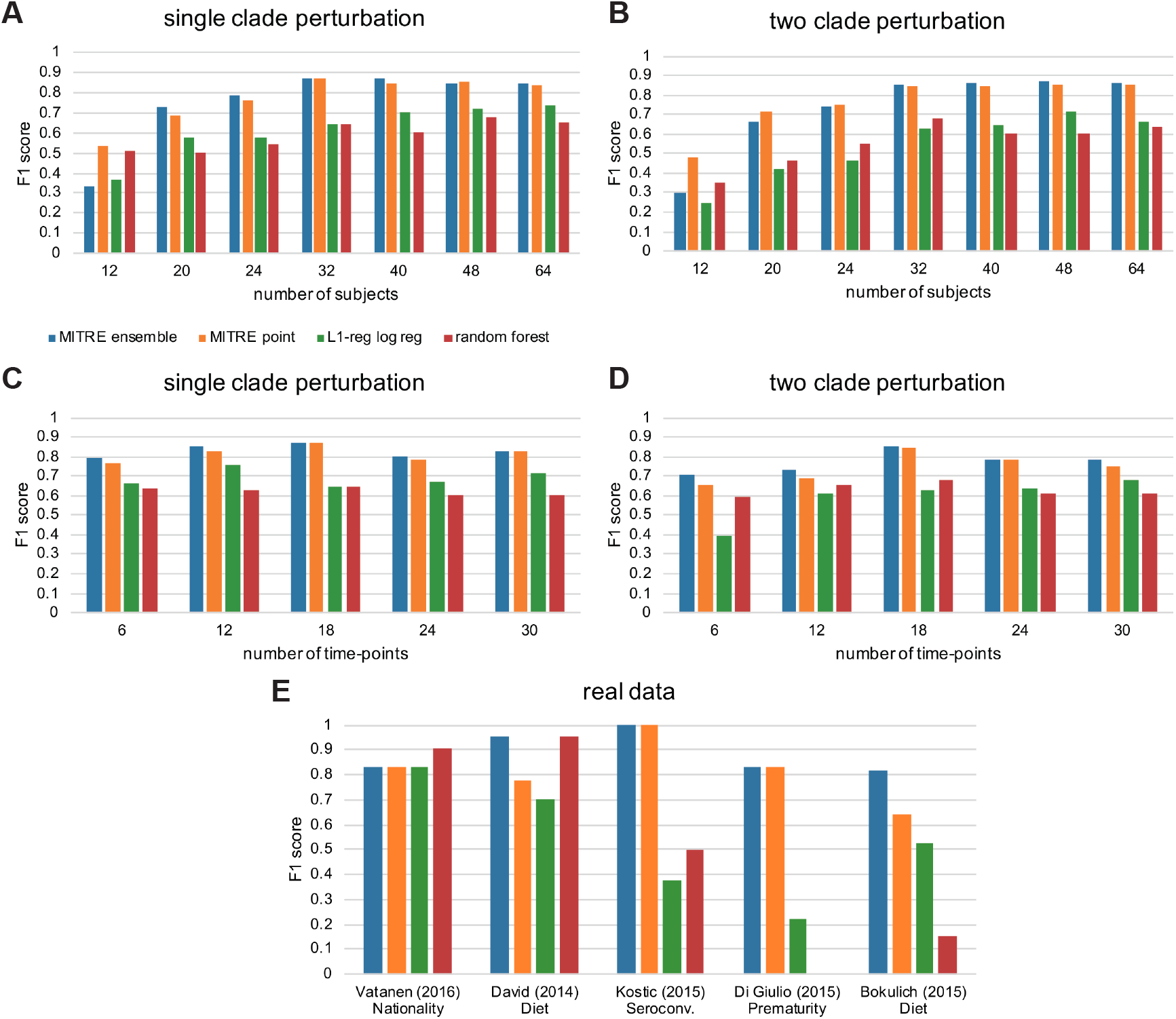
Cross-validated predictive performance of MITRE and comparator methods on semi-synthetic and real data. **(A-D)** Results on semi-synthetic data. A parametric bootstrapping-type method was used to generate simulated data from an underlying real dataset. Simulated cases were generated by randomly selecting and perturbing bacterial clades over a randomly selected limited time-window (∼20% of the duration of the study); an equal number of control subjects were simulated. For the one clade perturbation scenarios, the clade remained unperturbed for the simulated cases; for the two clade perturbation scenarios, one clade was perturbed in the simulated control subjects, and both were perturbed in the simulated cases. **(A-B)** One or two clades randomly perturbed in simulated subjects, 18 time-points, varying numbers of subjects. **(C-D)** One or two clades randomly perturbed in simulated subjects, 32 subjects, varying numbers of time-points. **(E)** Results on real data. The different methods were used to predict the indicated categories in the datasets shown. F1-score is the harmonic mean of precision and recall; higher scores indicate superior results.

### Benchmarking against standard machine learning methods: real data

We next evaluated the performance of MITRE on real experimental datasets with 16S rRNA amplicon and whole-genome-shotgun metagenomic sequencing data, from five representative published studies with sufficiently dense temporal sampling and numbers of subjects. Vatanen et al. [6] tracked gut microbiome composition and life history data, including allergy diagnoses and serum IgE levels, from birth to three years of age in cohorts of infants at high risk for autoimmune disease in Finland, Estonia, and Russia. David et al. [3] tracked the gut microbiome composition of healthy adults before, during and after a five-day period of consuming exclusively plant-based or exclusively animal-based diets. Bokulich et al. [2] tracked the gut microbiome composition from birth to two years of age in a cohort of U.S. infants and examined the effects of mode of delivery, diet, and antibiotic exposure. Kostic et al. [5] tracked the gut microbiome of Finnish and Estonian infants at high risk for type 1 diabetes throughout the first years of life, examining microbiome correlates of disease development. Finally, DiGiulio et al. tracked the composition of the vaginal microbiome in a cohort of pregnant women, investigating an association with premature delivery.

Using the microbiome and outcome/class data from these five studies, we defined a total of eleven representative microbiome-based prediction or classification tasks (e.g., given the vaginal microbiome data of DiGiulio et al., predict which women in the cohort experienced premature delivery; given the gut microbiome data of David et al, determine which time series correspond to exclusively animal-based diets vs exclusively plant-based diets; etc.) A full list of the tasks analyzed is given in Supplementary Table 1. In five out of the eleven tasks, we found that at least one of the methods performed well (F1-score > 0.7, indicating reasonably high precision and recall.) These tasks represent scenarios in which true biological signal may be present in the data, and thus serve as the most meaningful basis for comparing performance of the different methods. Detailed results for all methods on all tasks are given in Supplementary Table 1.

Both the MITRE point estimate and ensemble method achieved high accuracy on all five of the relevant prediction or classification tasks (Figure 2E). In the case of distinguishing infants fed formula-based diets from those predominantly breast-fed (Bokulich *et al*.), predicting seroconversion to serum autoantibody positivity in infants at high risk of T1D (Kostic *et al*.), and predicting premature delivery (DiGiulio *et al*.), MITRE significantly outperformed the random forest and L1-regularized logistic regression approaches; for Russian cohort membership (Vatanen *et al*.) or plant-based diet (David *et al*.) prediction, MITRE performed on par with the best comparator method (Random Forest). These results are consistent with our semi-synthetic data simulations as well. Collectively our results suggest that MITRE’s phylogenetic aggregation approach and robust use of the temporal structure of the data provide significant advantages for classification, and the increased interpretability of the MITRE point estimate comes at little if any cost in predictive accuracy.

### Model interpretability and exploratory analysis capabilities

We illustrate here an example demonstrating MITRE’s ability to achieve high accuracy while maintaining interpretability. The best (point estimate) rule set learned by MITRE to distinguish between predominantly formula-fed and predominantly breast-fed infants in the study of Bokulich et al. [2] is “If, between the 1^st^ day of life and the 156^th^ day of life, the average abundance of clade 13241 increases faster than 0.03% per day, the probability that the subject was predominantly formula-fed is 79%; otherwise that probability is 5.4%.” Though this MITRE rule set is simple enough to express in a single sentence, it outperforms Random Forest models that aggregate the predictions of over a thousand decision trees (Figure 2E). Moreover, this MITRE rule set lends itself readily to biological interpretation. Clade 13241 is a broad group of Firmicutes, including OTUs in the dataset mapping to *Ruminococcus gnavus*, *Roseburia hominis*, and several *Clostridium* and *Blautia* species. These species are generally viewed as more representative of adult or at least more mature microbiomes, being strict anaerobes with specialized carbon source utilization requirements and capabilities (e.g., [20]), suggesting that the formula diet may shift infants toward more adult-like gut microbiota. Of note, the expressiveness and interpretability of the MITRE rule set format is retained even in cases of nonlinear interactions across multiple clades and time windows; see the Supplementary Note for an example.

In addition to providing the point estimate, which can serve as a powerful predictive model as described, MITRE also allows the user to explore the distribution of probable rules learned by the framework. Such explorations can be useful for further interpreting rule sets and generating biological hypotheses. Figure 3 illustrates MITRE’s capabilities for interactive visualization of the distribution of learned rules. Heat maps as shown in Figure 3A and 3D allow the user to examine the time windows and regions of the phylogenetic tree where the temporal changes in the microbiota are most strongly associated with the outcome of interest. In Figure 3A-C, MITRE has been used to learn rules that distinguish the microbial dynamics observed when the subjects in the study of David et al. [3] were fed an exclusively plant-based diet for five days versus the dynamics observed when subjects were fed an exclusively animal-derived diet. The user has clicked on two areas on the heat map, revealing rules that apply to different time windows and different groups of OTUs in the order Clostridiales. The first rule set pertains to a clade containing *Roseburia* species, which are butyrate producers, whereas the second rule set pertains to a clade containing *Dorea* species, which produce other short-chain fatty acids including acetate and formate, but not butyrate. Thus, this capability to explore the distribution of rule sets allows the user to find evidence that the animal-based diet promotes two groups of phylogenetically distinct, and likely functionally distinct, groups of microbes.

**Figure 3.**
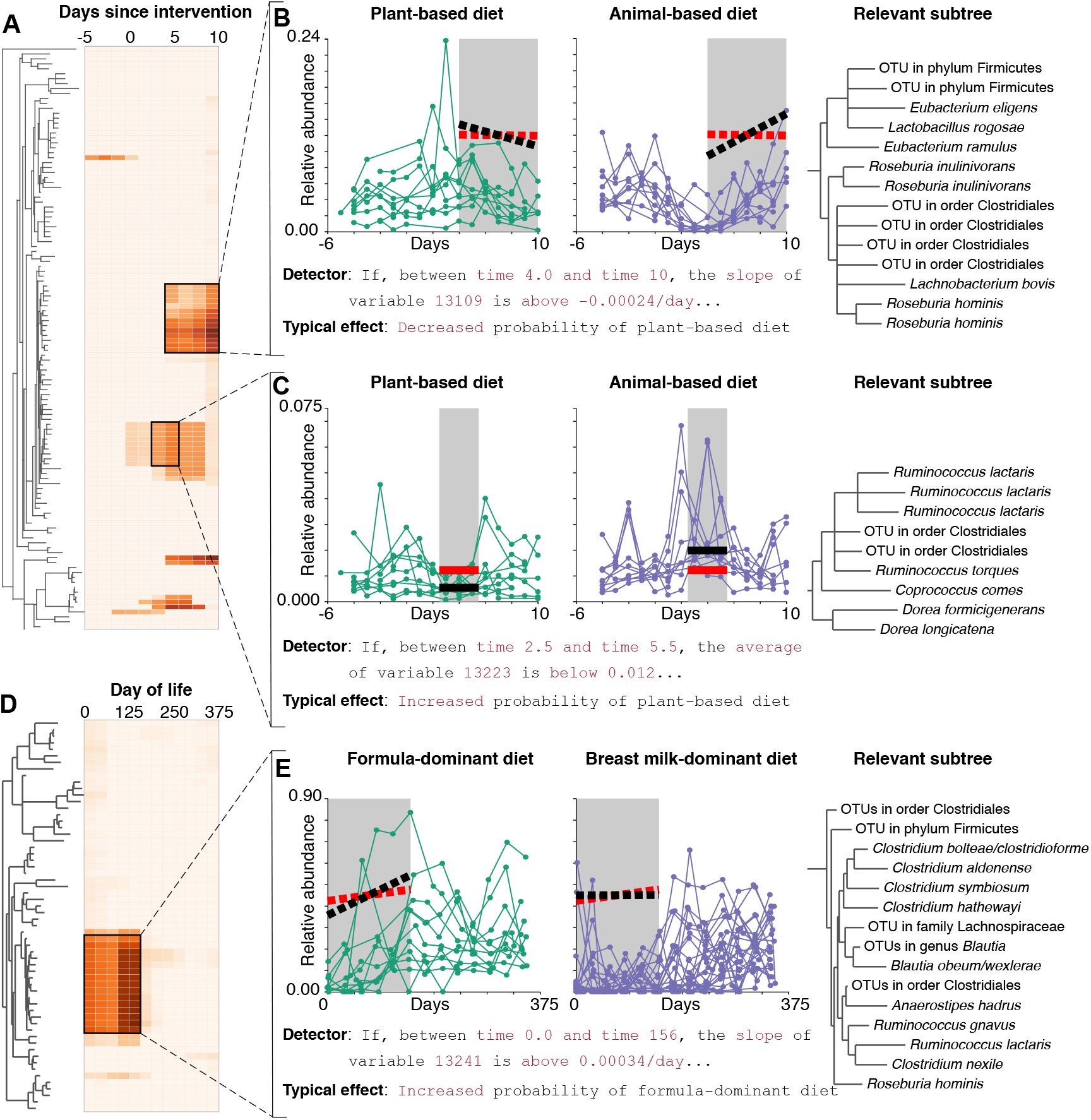
MITRE supports exploratory analyses through an interactive visualization interface. The interface allows the user to explore the distribution of learned rules. MITRE was applied to predict diet type from data from David et al. [3] **(A-C)** or Bokulich et al. [2] **(D-E)**. In **(A)** and **(D)**, cell colors indicate the strength of evidence that the dynamics of an OTU, or one of its ancestors, during a time window is associated with diet. Panels **(B)**, **(C)**, and **(E)** show high-probability detectors and phylogenetic subtrees to which they apply. **(B-C)** Analyses reveals dynamic behaviors of two different clades, one with butyrate producers and the other without, which distinguish subjects on plant or animal-based diets. The animal-based diet thus promotes two groups of phylogenetically distinct microbes which are also likely functionally distinct. **(E)** Analyses reveal dynamic behavior of a clade of bacteria, associated with a more mature microbiome, which is increased in the predominantly formula-fed infants, suggesting the formula diet may shift infants toward more adult-like gut microbiota. Red lines, threshold slopes/abundances; black lines, median slopes/abundances. Median effect = median over all rules containing the detector.

As another example, Figures 3D and 3E demonstrate exploration of the posterior distribution of rule sets identifying temporal patterns that distinguish the microbiota of predominantly formula-fed and predominantly breast-fed infants in the study of Bokulich et al. [2], for which the point estimate described above performed well. The heat map of Figure 3D shows that the high-posterior-probability rule sets are strongly focused on a single group of OTUs in the first 156 days of life. Figure 3E presents a particular detector, included in many such rule sets, that discriminates effectively between diet types in the training data (which is the detector used by MITRE to form the point estimate rule discussed above). Thus, in this case, the user finds evidence that the posterior distribution is essentially unimodal, with the point estimate alone characterizing temporal differences between the formula and breast-milk fed infants well.

## Discussion

MITRE offers a number of advantages over generic statistical or machine learning methods. Incorporation of phylogenetic information readily allows for biological interpretation of results, as discussed, whereas standard classification methods that evaluate each taxon independently clearly do not have this advantage. MITRE automatically learns time windows that are relevant to predicting host status, as opposed to generic approaches that require the user to manually specify periods of interest, which are generally unknown *a priori*. We have also highlighted the utility of MITRE’s human-readable rules. These rules can capture rich temporal patterns and nonlinear relationships among microbes, but remain interpretable, as they are composed of simple and understandable detectors. Indeed, as we have shown, MITRE rule sets are not only easy to understand, but a single MITRE rule can outperform black box machine learning methods that make predictions based on collections of hundreds to thousands of components, which are difficult to understand even individually.

Another important feature of the MITRE framework is that it is fully Bayesian. Bayesian models are increasingly being adopted in a variety of fields, in particular for biomedical applications (e.g., [21]), because they provide a unified framework that handles a number of key modeling and inferential issues, including incorporation of prior knowledge, accurate estimation of confidence in predictions, and principled comparisons of multiple models. Bayesian methods for model comparison have recently been highlighted as powerful alternatives [22] to traditional *p*-value based hypothesis testing [23], because Bayesian approaches allow direct comparison of multiple relevant alternative models, rather than just the ability to reject a null model that is often of little interest in itself. In particular, MITRE facilitates principled model comparisons by calculating Bayes factors [24], which quantify the strength of the evidence provided by the data for each of a set of competing models. We have also described how exploration of the posterior distribution of rule sets using the provided GUI in the MITRE software package allows the user to evaluate the possibility of multiple informative rule sets and formulate biological hypotheses about the data set.

Although MITRE as currently implemented is primarily designed to use taxonomic abundance profiles derived from 16S rRNA amplicon sequencing or WGS metagenomics data as input–currently the most common data types in longitudinal microbiome studies–the model and software can readily incorporate other time-series of additional data types, e.g., host physiological measurements, metabolomics, or metagenomics-derived functional profiles, as such data become more widely available. Other model features that could be readily added in the future include time-varying, continuous, and multiple host outcomes, e.g., multivariate host read-outs such as blood pressure, blood glucose, and body weight.

## Conclusion

We have demonstrated that our software package MITRE overcomes the unique challenges of predicting host outcomes from microbiome time-series data, while drawing on a well-established tradition of rule-based techniques in machine learning and artificial intelligence [25–27], and can perform as well as or better than “black box” machine learning methods while maintaining interpretability. This latter feature is critical as microbiome analyses move into clinical applications, in which patients and physicians necessarily place a premium on transparency and interpretability of results. We have provided an open source and user-friendly implementation of our method, which we expect will greatly aid investigators analyzing longitudinal host-microbiome studies and ultimately provide novel insights into the complex interplay between microbiome dynamics and host health and disease.

## Methods

### Operation of the software and input data requirements

The MITRE software is implemented in Python 2.7.3. MITRE and its dependencies are available through the Python Package Index, pypi.python.org, facilitating installation across multiple platforms. The software is run from the command-line, with parameters and other inputs specified using a straightforward configuration file format.

Each MITRE run requires four input files (for the standard case of 16S rRNA amplicon data): a table of OTU abundances in each sample, a table specifying the subject and timepoint associated with each sample, a table specifying the outcome (and optionally other data) associated with each subject, and phylogenetic placements of the OTUs on a reference tree. The user provides the three tables in comma-separated-value format, and phylogenetic placements in the .jplace format produced by pplacer. Alternative input data types, including taxonomic abundance profiles generated from WGS metagenomic data with Metaphlan, are described in the MITRE manual online.

The output of the software, described in detail below, includes textual summaries of a single best set of rules (the point estimate) and the distribution of probable alternative rule sets, as well as an HTML file providing a graphical interface for interactive visualization of the results.

### MITRE model details

#### Mathematical basis of the MITRE model

MITRE can be expressed as a hierarchical generative model that generates sets of rules of the form described above. The generative process, starting at the top of the hierarchy, is as follows:

- Sample a length *K* for the rule set
- For each rule *k* ∈ {1,…,*K*}, sample:
  - A weight *β_k_*
  - A rule length *L_k_* (number of detectors in the rule)
  - The detectors *d* ∈ {1,…,*L_k_*} drawing from a pre-defined pool of detectors (see below) according to a probability distribution which is parameterized by the time window and bacterial group (phylogenetic subtrees) to which each detector applies.

The rules are then weighted in a Bayesian logistic regression model to predict host status. To be precise, assume we have observations *x_ijt_* for *i*=1,2,…,*N_subjects_* of relative abundances of bacterial OTUs *j*=1,2,…,*N*_OTUs_ sampled at time-points *t*=1,2,…,*N*_samples,*i*_, as well as a binary status variable *y_i_* for each host. The MITRE probability model can then be expressed as:

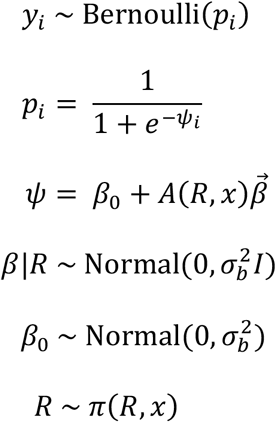

Here, *R* is a set of rules, and *A(R,x)* is the matrix whose entry *A_ik_* is 1 if the data from subject *i* satisfies the conditions of all detectors in the *k*-th rule in the set, and zero otherwise. Conditional on *R*, this is a standard Bayesian logistic regression model, whose covariates are the truth values of the rules in *R* (along with an offset term modeling baseline odds). MITRE also allows the optional inclusion of static non-microbiome covariates; see the Supplemental Note for complete details.

The prior probability distribution over rule sets, *π(R,x)*, is a mixture over the probability *Θ*_empty_ for an empty rule set *R_0,_* versus a truncated negative binomial distribution for the length of a non-empty rule set. For a non-empty rule set, the prior further models the distribution over detectors that comprise each rule, taking into account the length of the time window for the detector and an associated position on the phylogenetic tree. Hyperparameters for these priors, as well as priors on other variables, complete the model. A full specification of priors in MITRE, and a discussion of sensitivity to the choice of hyperparameters, are given in the Supplementary Note.

### Generation of pools of detectors from data

MITRE generates a comprehensive pool of detectors from the supplied data and user-specified parameters *t*_min_, *t*_max_, *N*_w_, and *N*_θ_, as follows:

1. Divide the duration of the experiment/observations into *N_w_* equal basic time windows, and enumerate all combinations of 1 or more consecutive basic time windows that are longer than *t*_min_ and shorter than *t*_max_ and during which at least one sample was collected for every subject.
2. Within each such time window *(t_0_, t_1_)*, for each bacterial group (phylogenetic subtree) *j*, calculate the average abundance of the group in each subject *i*, and sort those values, obtaining *v_1_* ≤ *v_2_* ≤ … ≤ *v_N_*_subjects_. If *N*_subjects_ < *N*_θ_+1, for *l*=1,2,…,*N*_subjects_-1, let *θ_l_* = (*v_l_* + *v_l+1_*)/2 and add the detectors “between *t_0_* and *t_1_*, the average abundance of group *j* is above *θ_l_* and “between *t_0_* and *t_1_*, the average abundance of group *j* is below *θ_l_*” to the population. If instead *N*_subjects_ > *N*_θ_, hierarchically cluster the values *v_0_*, …, *v_N_*_subjects_ into *N*_θ_ groups, let *θ_l_* be the midpoint between cluster *l* and cluster *l+1,* for *l*=1,2,…, *N*_θ_ −1, and add the detectors corresponding to those thresholds instead.
3. Then repeat the process for all combinations of 1 or more consecutive basic time windows longer than *t*_min_ and shorter than *t*_max_ during which at least two samples were collected for every subject, calculating the slope of the abundance of each group *j* in each subject *i* during each such window, and adding the detectors “If, between *t_0_* and *t_1_*, the slope of the abundance of group *j* is [above/below] *θ _l_*” to the population (again carrying out a clustering process to reduce the number of threshold values to *N* _θ_, if needed.)

The runtime of the inference algorithm (described below) depends approximately linearly on the size of the detector pool; thus the choice of parameters *t*_min_, *t*_max_, *N*_w_, and *N*_θ_ controls a tradeoff between high resolution (temporally, and in the space of threshold values for OTU abundance/slope) and performance. It is recommended to choose *N_w_* as large as possible while ensuring that most basic time windows include at least one observation from every subject, and to set *N*_θ_= 40; t_min_ should generally be set to 0 and t_max_ to either the duration of the study or (to enforce temporal localization of the rules in cases where, e.g., dramatic increases in abundance at a well-defined time also lead to notable increases in average abundance over the period of the entire study) half that duration, but may be adjusted if dynamics on a particular time scale are of *a priori* interest.

### Model inference

We perform approximate Bayesian inference, to learn the posterior distribution over model parameters including rule sets *R* and regression coefficients *β.* MITRE employs a custom Markov Chain Monte Carlo (MCMC) algorithm, which alternates efficient updates of the regression coefficients using a Polya-Gamma auxiliary variable scheme [28], Metropolis-Hastings update steps that propose changes to the rule set *R,* and updates to the hyperparameters governing the prior distribution over rule sets. The MCMC algorithm is described in detail in the Supplementary Note. Briefly, four types of updates to *R* are considered:

1. A randomly-chosen detector in *R* may be replaced by another detector from the pool
2. A randomly-chosen detector in *R* may be removed from *R* (if it is the only detector in a rule, the rule is removed as well)
3. A detector from the pool may be added to *R*, either to an existing rule chosen at random, or to form a new rule of length 1
4. A detector may be moved from one rule in *R* to another

For the analyses presented here, 50,000 samples were drawn from the posterior distribution (except for analyses of data from Vatanen et al. [6], where 25,000 samples were used, and Kostic et al. [5], where 100,000 samples were used.) Mixing of the MCMC sampler was assessed using the diagnostics described in the Supplementary Note; we recommend users employ these diagnostics to determine the appropriate number of samples needed for their particular studies.

Run time depends on the size and complexity of the data set; using a single Intel Xeon CPU (E5-2697 v3 2.60GHz), sampling took 45 minutes for the data of DiGiulio et al. [7], 23 hours for the data of David et al. [3], 30 hours for the data of Bokulich et al. [2], and 64.5 hours for the data of Vatanen et al. [6]. Cross-validation was performed in parallel (1 fold/core) requiring similar total elapsed time for each study.

### Data simulation

We simulated from the Bokulich et al. [2] dataset using a parametric bootstrapping-style procedure. Simulated subjects were sampled with replacement from the set of control subjects in the real data. Equal numbers of simulated cases and control subjects were generated for each scenario. In order to have sufficient real data to bootstrap to evaluate the range of scenarios of interest, we excluded subjects with fewer than 13 time-points sampled, who had no samples before 10 days, and who were studied for less than 600 days; this yielded 20 subjects for bootstrapping. We then truncated data to an interval between 10 and 600 days, since this contained the densest sampling across subjects.

Perturbations of duration 120 days, in time windows randomly occurring throughout the experiment, were introduced into randomly selected clade(s) to simulate cases, with the magnitude of the perturbation distributed among clade members according to the relative abundance of members in the original data. The duration of the perturbation(s) was chosen to be approximately of the order of that seen with the MITRE point rule on the real data set. Perturbations were introduced randomly into clades with the following characteristics: minimum average relative abundance of 0.1%, maximum average relative abundance of 20%, and a maximum of 30 OTUs in the clade. These parameters were chosen so as to provide a meaningful relative disturbance to other clades, but not to drastically disrupt the entire microbiome (which would present less of a challenge to the prediction algorithms.) The magnitude of perturbation(s) were sampled for each subject from log-normal distributions, with mean and variance of the order of that seen with the MITRE point rule on the real data set (control log mean = −6, control std = 1.5, case log mean = −3, case log std = 1.5). When two perturbations were applied, each control subject received only one perturbation, whereas case subjects received both perturbations.

Note that MITRE is a supervised learning (conditional) method, meaning that the microbiome data itself is not modeled; to simulate time-points not present in the original data set, we therefore must introduce a model of microbiome dynamics. We model the underlying microbiome data as arising from latent time-dependent stochastic processes (Gaussian random walks):

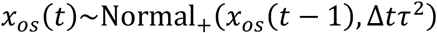

Here, *x_os_(t)* is the latent trajectory for OTU *o* in subject *s* at time *t*, and *τ^2^* is the process variance parameter, which is empirically estimated from the real data as approximately the 75-percentile variance. We assume a Bayesian model and infer the posterior latent trajectories using a 1-step ahead MCMC algorithm similar to our previously described method [29], except in this case trajectories are assumed to be independent of one another.

The observed data *c_s_(t)*, consisting of sequencing counts, is assumed to arise through a two-stage error model:

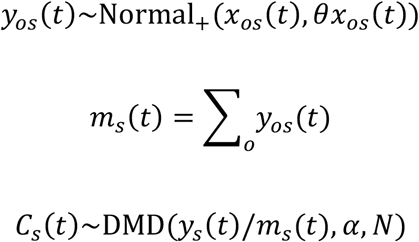

Here, DMD denotes the Dirichlet-Multinomial distribution with concentration parameter *α* and number of simulated sequencing reads per sample *N*; we use parameters estimated from data (*a* = 286; *N* = 50,000) as previously described [14].

The model thus provides temporal coherence through the Gaussian random walk latent trajectory, while modeling compositionality and over-inflation through the two-stage error model. Posterior samples from the model capture temporal trends seen in the real data (e.g., periods of time in which a particular OTU are increasing), but with randomness introduced so that subjects sampled with replacement look sufficiently different.

For each scenario simulated (e.g., a particular number of time-points or subjects), 10 independent simulations were performed. To investigate the effects of varying the number of subjects, we fixed the simulation at an intermediate number of time-points (18), and simulated different numbers of subjects. Similarly, to investigate the effects of varying the number of time-points, we fixed the simulation at an intermediate number of subjects (32), and simulated different numbers of time-points. To facilitate comparisons, we sampled evenly spaced time-points in all cases. The simulated data were then provided to MITRE and the other methods in the same input format as for real data, as described below. Complete Python code to reproduce the simulations is available in the MITRE online repository.

### Output

MITRE generates several summaries from the posterior samples obtained from the MCMC inference procedure described above. The *point estimate* is a single rule set *R\** (with coefficients *β\**) that summarizes the mode of the posterior distribution. If the posterior probability that *R* is empty is greater than 0.5, the point estimate is the empty rule set. Otherwise, to obtain a representative non-empty summary list, we determine the posterior mode *d\** of the total number of detectors in *R*, and take *R\** to be the rule set with the highest posterior probability among all sampled rule sets that contain *d\** total detectors. The point estimate coefficients *β\** are the highest posterior probability sampled coefficients associated with *R*.*

To provide an overview of the possible alternative rule sets learned by the model, rule sets in the posterior samples are clustered and a summary of the highest-probability clusters is produced. The clustering process first forms clusters of detectors that apply to the average value or slope of the same variable in highly overlapping time windows (ignoring threshold values), then clusters together rules whose component detectors belong to the same clusters, and finally groups rule sets whose rules belong to the same clusters (see the Supplementary Note for a full description.) For each cluster, a representative rule set and the estimated posterior probability that *R\** belongs to the cluster are reported.

Calculation of the Bayes factor for the empty rule set *R_0_* (vs any non-empty *R*) and two additional types of Bayes factors, indicating the relevance of phylogenetic subtrees or time-windows, is described in the Supplementary Note.

Finally, MITRE generates an interactive graphical visualization of the posterior distribution of rule sets. A heatmap of the Bayes factors for leaf variable/basic time window combinations is rendered alongside the bacterial phylogeny (as in figure 3A, 3D); clicking on any cell allows the visualization of the detectors associated with the cell with the largest Bayes factors (as in figure 3B-C and 3E.)

### Data preprocessing and filtering

MITRE offers a number of user-configurable options for preprocessing and filtering microbiome time-series data. The following procedure is recommended, and was used for the results presented here (except as noted below):

1 To remove potentially spurious rare OTUs, discard all OTUs with less than *N*_counts,OTU_ observed across all samples (typically *N*_counts,OTU_ = 10.)
2 To exclude samples where coverage is so low that abundance estimates for uncommon OTUs may be unreliable, discard all samples with less than *N*_counts,sample_ total counts observed across all remaining OTUs (typically *N*_counts,sample_=5000 for HiSeq/MiSeq data.)
3 If desired, to analyze only a particular time period (because, e.g., samples are not available outside that period for the majority of subjects) discard all samples before time *t_i_* and after time *t_f_* (by default, *t_i_* is the time of the earliest available sample, and *t_f_* the time of the latest sample.)
4 To exclude subjects for whom microbiome dynamics cannot be resolved at the desired temporal resolution throughout the entire study, discard subjects with too few, or too sparse, observation points, by dividing the duration of the study into *N*_w,filter_ equal pieces and keeping only subjects with at least *N_s_* samples in any *N_c_* consecutive such pieces. Default values are *N*_w,filter_=10, *N_s_ =* 2, and *N_c_*=1; note that for data with very inconsistent sampling, these parameters must be chosen judiciously to maximize the number of subjects included while allowing an acceptable level of temporal resolution.

After these steps, counts data are converted to relative abundance data for each sample, and, for each node in the phylogenetic tree, a relative abundance estimate is obtained by summing the relative abundances of its children. The following filtering steps are then applied to all taxa (both OTUs and higher nodes in the tree):

5 To exclude low abundance taxa (for which abundance estimates may be inaccurate) or infrequently observed taxa (which we expect are unlikely to be explanatory, though higher taxa including them may be), discard all taxa except those that exceed a threshold abundance *a* in at least *N_a_* consecutive samples in at least *N_i_* subjects. Typically, *N_a_=*2. Appropriate values for *a* and *N_i_* depend on the number of subjects and typical reads per sample; for studies with 10-150 subjects and average reads per sample on the order of 10^4^, we recommend *a=*10^-4^ and N_i_=4 or 10% of the number of subjects, whichever is larger.
6 To minimize redundancy, discard all taxa corresponding to nodes in the phylogenetic tree with exactly one remaining child taxon, as their temporal patterns are often very similar to those of their children.

Note that, when a large number of taxa are considered, the detector pool becomes large and the computational cost of the inference algorithm grows; if necessary, it is recommended to increase the stringency of steps 5 and 6 to keep the total number of taxa below 500.

### Bioinformatics and preprocessing for analyzed datasets

For each 16S-based dataset to which MITRE was applied, the original 16S rRNA amplicon sequencing data was reprocessed to obtain tables of OTU abundances and phylogenetic placements for each OTU on a reference tree, using as consistent an analysis process as possible given differences in sequencing methodology and the nature of the available data. Where possible, DADA2 1.1.5 [30] was used to trim, quality filter, merge, and remove chimeras from the reads, assign them to inferred true sequences, and classify the inferred sequences taxonomically (such classification is not necessary for MITRE application, but is helpful for interpretation of the results.) Inferred sequences were then placed on a reference tree generated from full-length or near full-length (>1200 nt) 16S rDNA sequences of type strains from the Ribosomal Database Project [31] using pplacer [18]. When quality scores for sequences were not available and DADA2 could not be used, sequences were instead processed using mothur 1.35.1 [32, 33] for denoising, quality filtering, alignment against the ARB Silva 16S gene sequence reference database, clustering into OTUs at 97% identity, and taxonomic classification. For the WGS metagenomics data of Kostic et al [5], published taxonomic abundance tables were used directly as input data to MITRE, exploiting MITRE’s built-in support for parsing Metaphlan result tables, described in the MITRE manual. Full details regarding pre-processing of all datasets are described in the Supplemental Note.

After reanalyzing sequencing data for each study and excluding subjects with infrequent sampling (see description of filtering methods above) we applied MITRE and the comparator methods to classify subjects according to relevant categories: membership in the Russian cohort (n=30), elevated IgE levels (n=28), diagnosis with any allergy (n=49), any dietary allergy (n=42), with dairy allergy (n=32), or with egg allergy (n=25) in the data of Vatanen et al. (n=113 total for nationality; n=109 total for all other outcomes); formula-dominant diet (n=11) or Cesarean delivery (n=13) in the data of Bokulich et al. (n=35 total); seroconversion (n=11) in the data of Kostic et al. (n=19 total); premature delivery (n=6) in the data of DiGiulio et al. (n=37 total); and plant-based diet (n=10) in the data of David et al. (n=20 total.)

### Comparison methods

To compare MITRE’s predictive performance to alternative methods, each OTU’s abundance data for each subject was averaged across all observations within each of a set of time windows, defined by dividing the experiment into *N*_w,comparison_ equal intervals and taking any consecutive *N*_c,comparison_ such intervals as a valid time window. Parameters were chosen to maximize temporal resolution while ensuring that each time window still contained at least one observation for each subject.

Note that the same subjects were used (i.e., those not excluded by the preprocessing settings described above) for both MITRE and the comparison methods. For David et al. [3], *N*_w,comparison_=10 and *N*_c,comparison_=3; for Vatanen et al. [6], *N*_w,comparison_=9 and *N*_c,comparison_=4; for DiGiulio et al. [7], *N*_w,comparison_=*N*_c,comparison_=1; for Bokulich et al. [2], *N*_w,comparison_=12 and *N*_c,comparison_=2; for Kostic et al. [5], *N*_w,comparison_=5 and *N*_c,comparison_=2.

These averaged abundances were then used to train Random Forest or logistic regression classifiers using the Python package scikit-learn [34]. Random Forest classifiers included 1024 trees (as larger numbers of trees were not found to improve classifier performance.) For logistic regression with L1 regularization, the regularization strength parameter was chosen using 10-fold crossvalidation from among a grid of logarithmically spaced options spanning the range 10^-4^ to 10^4^.

## Supporting information

Supplemental Details

## Declarations

### Acknowledgements

We thank Daniel DiGiulio for assistance with the data from reference [7], and Travis Gibson for helpful comments on the manuscript.

### Availability of data and materials

The MITRE software package is available at: https://github.com/gerberlab/mitre/. All additional input files needed to reproduce the results presented here, and detailed output from all MITRE simulations discussed, are available at https://github.com/gerberlab/mitre_paper_results.

Datasets analyzed during the current study are available: data from Bokulich et al. [2], from ENA (https://www.ebi.ac.uk/ena/data/view/PRJEB14529) and QIITA (https://qiita.ucsd.edu, study 10249); data from David et al. [3], from MG-RAST (http://metagenomics.anl.gov/linkin.cgi?project=mgp6248); data from Kostic et al.

[5], via the DIABIMMUNE project web site (https://pubs.broadinstitute.org/diabimmune/t1d-cohort); data from Vatanen et al.

[6], via the DIABIMMUNE project web site (https://pubs.broadinstitute.org/diabimmune/three-country-cohort); data from DiGiulio et al. [7], on request from those authors.

### Authors’ contributions

EB and GKG conceived the method, developed the theory, and wrote the manuscript. EB wrote the software, curated the relevant data, and formally analyzed and validated the method and software. RC contributed to the software and analysis and validation of the method.

### Ethics approval and consent to participate

Not applicable

### Consent for publication

Not applicable

