## Supplemental Table 1 for "MITRE: predicting host status from microbiota time-series data"

| Data from study | Variable predicted | MITRE ensemble F1 | MITRE point F1 | RF F1 | LR F1 | N, 'positive' outcome | N, total | CV type |
|---|---|---|---|---|---|---|---|---|
| David *et al.* (2014) | Plant-based vs. animal-based diet | 0.952 | 0.778 | 0.952 | 0.7 | 10 | 20 | leave-one-out |
| Kostic *et al.* (2015) | Seroconversion | 1.0 | 1.0 | 0.5 | 0.375 | 11 | 19 | leave-one-out |
| Di Giulio *et al.* (2015) | Premature delivery | 0.833 | 0.833 | 0.0 | 0.222 | 6 | 37 | leave-one-out |
| Bokulich *et al.* (2016) | Formula-dominant diet | 0.818 | 0.64 | 0.154 | 0.526 | 11 | 35 | leave-one-out |
| Bokulich *et al.* (2016) | Cesarean delivery | 0.211 | 0.643 | 0.3 | 0.0 | 13 | 35 | leave-one-out |
| Vatanen *et al.* (2016) | Russian nationality | 0.833 | 0.833 | 0.909 | 0.833 | 30 | 113 | 5-fold |
| Vatanen *et al.* (2016) | Any allergy | 0.353 | 0.621 | 0.556 | 0.4 | 49 | 109 | 5-fold |
| Vatanen *et al.* (2016) | Any dietary allergy | 0.0 | 0.0 | 0.2 | 0.0 | 42 | 109 | 5-fold |
| Vatanen *et al.* (2016) | Egg allergy | 0.0 | 0.0 | 0.0 | 0.0 | 25 | 109 | 5-fold |
| Vatanen *et al.* (2016) | Dairy allergy | 0.0 | 0.0 | 0.0 | 0.0 | 32 | 109 | 5-fold |
| Vatanen *et al.* (2016) | Elevated IgE levels | 0.0 | 0.0 | 0.0 | 0.0 | 28 | 109 | 5-fold |

**Supplementary Table 1.** The classification problems to which MITRE and the comparator methods were applied, and the performance of the methods applied to each. Full references are given in the main text. All F1 scores shown are results of cross-validation of the type shown. RF, random forest; LR, L1-regularized logistic regression; N, number of subjects; CV, crossvalidation. Total number of subjects counts only those remaining after data filtering and preprocessing as described in the manuscript and Supplementary Note (and discarding subjects for which data about the variable of interest was not available). Note that the total number of subjects given for the study of David *et al* represents the total number of time series available for classification; most individuals received plant-based and animal-based diets in sequence (with washout periods), so the true number of subjects is less than 20.
