## Supplemental Table 2 for "MITRE: predicting host status from microbiota time-series data"

| Data from study | Variable predicted | max $\hat{R}$ | iterations | comparator features | pool |
|---|---|---|---|---|---|
| David *et al.* (2014) | Plant-based vs. animal-based diet | 1.014 | 50000 | 66184; 1344 | 374848 |
| David *et al.* (2014) | Diet (data renormalized) | 1.011 | 50000 | 66184; 2096 | 519954 |
| Kostic *et al.* (2015) | Seroconversion | 1.0060 | 100000 | 1520; 224 | 12278 |
| Di Giulio *et al.* (2015) | Premature delivery | 1.033 | 50000 | 1551; 23 | 3516 |
| Bokulich *et al.* (2016) | Formula-dominant diet | 1.028 | 50000 | 25344; 528 | 397884 |
| Bokulich *et al.* (2016) | Cesarean delivery | 1.031 | 50000 | 25344; 528 | 397884 |
| Vatanen *et al.* (2016) | Russian nationality | 1.043 | 25000 | 18180; 1248 | 570934 |
| Vatanen *et al.* (2016) | Any allergy | 1.036 | 25000 | 17646; 1212 | 555896 |
| Vatanen *et al.* (2016) | Any dietary allergy | 1.024 | 25000 | 17646; 1212 | 555896 |
| Vatanen *et al.* (2016) | Egg allergy | 1.061 | 25000 | 17646; 1212 | 555896 |
| Vatanen *et al.* (2016) | Dairy allergy | 1.068 | 25000 | 17646; 1212 | 555896 |
| Vatanen *et al.* (2016) | Elevated IgE levels | 1.032 | 25000 | 17646; 1212 | 555896 |

**Supplementary table 2.** Technical details of the application of MITRE and comparator methods to each classification problem listed in supplementary table 1, plus the dietary classification problem applied to the data of David *et al.* normalized relative to the abundance of the genus Bacteroides, as shown in Supplementary Figure 1 and discussed in the text. R-hat values are calculated using five independent MCMC chains of the indicated length and represent the minimum value across the parameters monitored (i.e., parameters in the model, except that some high-dimensional vector-valued parameters were downsampled.) 'Pool' is the MITRE detector pool size. The comparator methods were applied both before and after the reduction of the number of variables by the abundance filtering options described in the text and Supplementary Note (reporting the results from whichever choice led to better performance for each method); the applications at these steps led to the larger and smaller numbers of features shown.
