## Supplemental Methods for "MITRE: predicting host status from microbiota time-series data"

**6 An example rule set with interactions inferred from real data 26**

### 1 MITRE model description

#### 1.1 Overview

MITRE uses a hierarchical, fully-Bayesian generative model. The model generates sets of rules $R$, with each rule consisting of a conjunction of *detectors*. The rules are then weighted in a Bayesian linear regression model to predict host status $y_s$.

#### 1.2 Inputs

As input to the model, assume that we have measurements $x_{os}(t_{s1}), x_{os}(t_{s2}), \ldots$ of the abundances of OTUs $o \in \{1, \ldots, n_{\text{OTUs}}\}$ in samples from subjects $s \in \{1, \ldots, n_{\text{subjects}}\}$.

Observation timepoints $t_{s1}, t_{s2}, \ldots$ may vary across subjects, but are assumed to all occur in the window $[0, T_{\text{experiment}}]$. We denote the total number of observations for subject $s$ as $N_s$ and write $x_{osk} = x_{os}(t_{sk})$. For each subject we observe a binary host status variable $y_s$, which we assume is time-independent.

We also assume as input a phylogeny of the OTUs as a tree with $n_{\text{clades}}$ nodes. We describe the set of OTUs descended from node $i$ as clade $i$, with the clades associated with the leaves of the tree being the OTUs themselves. Section 3.3 addresses the construction and interpretation of such trees in greater detail.

#### 1.3 Detectors and rules

Rules in MITRE consist of conjunctions of *detectors*, which are statements taking on values of 0 (false) or 1 (true). A MITRE detector $r$ is of the form:

> Between time $t_0$ and time $t_1$, the [average value/slope] of the abundance of clade $i$ is [above/below] $\theta$.

We refer to the parameters defining a detector as its *variable* ($i$), *time window* $[t_0, t_1]$, *type* ("average" or "slope"), *direction* ("above" or "below") and *threshold* ($\theta$).

Let $\mathbf{v}(r)$ denote the $n_{\text{subjects}}$-dimensional vector with entries $\mathbf{v}(r)_i = 1$ if the detector is true for subject $i$ and 0 otherwise.

A MITRE rule $\rho$ can thus be written as:

$$(r_{j1}, \ldots, r_{jn_1})$$

The value of rule $\rho$ for subject $i$ is thus $\prod_j \mathbf{v}(r_j)_i$, i.e., its value is 1 if all its detectors are true, and 0 otherwise.

#### 1.4 Detector population

In principle, the parameters for detectors $\theta$, $t_0$, and $t_1$ could be treated as continuous. However, to make inference efficient, we discretize parameter values and enumerate a finite population $\mathcal{D}$ of possible detectors as described below.

**Discretized time windows** We divide the experiment into $N_w$ equal time intervals

$$[0, (1/N_w)T_{\text{experiment}}], [(1/N_w)T_{\text{experiment}}, (2/N_w)T_{\text{experiment}}] \ldots.$$

A detector time window may be any consecutive combination of one or more of these time intervals, subject to the restrictions that the total length of the combination must be at least a minimum length $W_{\min}$ (user-specified) and no longer than a pre-defined maximum length $W_{\max}$ (user-specified, with a default value equal to the duration of the exoperiment,) and that detectors applying to average values may apply only over time windows within which all subjects have at least one associated observation, and detectors applying to slope values may apply only over time windows within which all subjects have at least two associated observations. Thus, when $N_w$ is sufficiently large that not all intervals contain at least one observation from all subjects, detectors with higher temporal resolution (shorter windows) are adaptively generated during times for which more samples are available. Note there are at most $N_w(N_w + 1)/2$ valid time windows.

**Discretized threshold values** To determine the set of possible threshold values for a detector with variable $i$ and time window $[t_0, t_1]$, we evaluate the average value or slope (as appropriate) of variable $i$ in that time window for each subject $s$. The average value of variable $i$ for subject $s$ in the time window $[t_0, t_1]$ is estimated in the natural way: it is the mean value of $x_{isk}$ over all data points for subject $s$ such that $t_0 \leq t_{sk} \leq t_1$.

To estimate the slope, we change variables to $\tau = t - (t_0 + t_1)/2$ and use the data points for that subject within the time window to fit a linear model $x_{is} = \alpha_{is}\tau + \beta_{is}$, taking the ordinary least-squares estimate of $\alpha_{is}$ as the slope.

(Note that detectors are not allowed that would require us to evaluate a slope over a window in which fewer than two observations are available, or an average over a window in which no data is available.)

Denote the results of the average and slope calculations, ordered from smallest to largest, $\zeta_1, \ldots, \zeta_{n_{\text{subjects}}}$, and denote the (possibly smaller set) of unique results as $\zeta_1 < \ldots < \zeta_{n_{\text{unique}}}$. A maximum number of threshold values per variable-time window-type-direction combination, $N_\theta$, is set by the user.

Two cases are considered. If $n_{\text{unique}} \leq N_\theta + 1$, then the allowed threshold values are set as the midpoints between the observed values, $(\zeta_1 + \zeta_2)/2$, $(\zeta_2 + \zeta_3)/2$, etc. Otherwise, a hierarchical clustering of the values $d_1, \ldots, d_{n_{\text{unique}}}$ is formed using the UPGMA algorithm[1], and truncated to produce a flat clustering of the $\zeta$ values into $N_\theta + 1$ groups; denote these $Z_1, \ldots, Z_{N_\theta+1}$, ordering them so that all values in $Z_i$ are less than all values in $Z_{j>i}$. The allowed threshold values are then the $N_\theta$ *midpoints of the gaps between clusters*, $(\max(Z_1) + \min(Z_2))/2, (\max(Z_2) + \min(Z_3))/2$, etc.

With this specification, the total number of detectors $n_{\mathcal{R}}$ will be at most

$$n_{\mathcal{R},\max} = (\text{number of clades after preprocessing}) \cdot N_w(N_w + 1)/2 \cdot N_\theta \cdot 4,$$

the factor of 4 arising from the possible combinations of type (average value vs slope) and direction. Note that, by construction, no detector will be true for all subjects or false for all subjects.

[1] R. Sokal and C.D. Michener, *University of Kansas Science Bulletin* 38, 1409-1438 (1958).

#### 1.5 Prediction of host status

MITRE predicts host status $y_i$ from sets of rules using a Bayesian logistic regression model defined as follows:

$$
\begin{aligned}
y_i &\sim \text{Bernoulli}(p_i) \\
p_i &= \frac{1}{1+e^{-\psi_i}} \\
\boldsymbol{\psi} &= \beta_0 + A(R,x)\boldsymbol{\beta} \\
\boldsymbol{\beta}|R &\sim \text{N}(\mathbf{0}, \sigma_b^2 I) \\
\beta_0 &\sim \text{N}(\mathbf{0}, \sigma_b^2)
\end{aligned}
\tag{1}
$$

Here, $R$ is a (possibly empty) ordered collection of (nonempty) collections of detectors from the pool $\mathcal{D}$ defined above:

$$
R = \left( \underbrace{(r_{11}, \ldots, r_{1n_1})}_{\rho_1}, \ldots \underbrace{(r_{m1}, \ldots, r_{mn_m})}_{\rho_m} \right).
$$

The matrix $A(R,x)$ is obtained from $R$ as:

$$
\left( \; \mathbf{v}_0 \;\middle|\; \mathbf{v}(r_{11}) \cdot \mathbf{v}(r_{12}) \cdot \ldots \cdot \mathbf{v}(r_{1n_1}) \;\middle|\; \cdots \;\middle|\; \mathbf{v}(r_{m1}) \cdot \mathbf{v}(r_{m2}) \cdot \ldots \cdot \mathbf{v}(r_{mn_m}) \; \right)
$$

where $v_0 = (1, 1, \ldots, 1)^T$. The remaining columns of $A$ indicate whether each rule in $R$ is true or false for each subject, and are obtained by taking the element-wise products of the vectors giving the truth of each detector within each rule. For example, if we have three subjects, $R = ((r_{11}, r_{12}), (r_{21}))$, and $\mathbf{v}(r_{11}) = (1, 1, 0)$, $\mathbf{v}(r_{11}) = (1, 0, 1)$, and $\mathbf{v}(r_{11}) = (0, 1, 0)$, then

$$
A(R) = \begin{pmatrix} 1 & 1 & 0 \\ 1 & 0 & 1 \\ 1 & 0 & 0 \end{pmatrix}.
$$

(In the interests of concision, we abbreviate $A(R,x)$ as $A(R)$ below, but it always implicitly depends on the data $x$ through the vectors of truth values $\mathbf{v}(r_{ij})$.)

Note that the dimension of $R$ is unknown; it is learned as part of the model. Thus, the prior probability distribution on $\boldsymbol{\beta}$ is conditional on $R$ because the length of $R$ determines the dimensions of $A$ and thus of $\boldsymbol{\beta}$.

**Comment on ordering of rules and detectors** For convenience, we have presented $R$ such that the detectors $r_{i1}, r_{i2}, \ldots, r_{in_i}$ within each rule $\rho_i$ are ordered, but it is clear that $A(R)$ and thus the model likelihood do not depend on this ordering. In actuality, we formally handle rules as unordered collections (sets). Thus, $\rho_i$ is formally a vector $z^{(i)} \in \mathbb{Z}_{\geq 0}^{n_\mathcal{D}}$ whose $j$th element $z_j^{(i)}$ gives the number of copies of detector $j$ included in the rule $\rho_i$. (Multiple copies of the same detector within a rule are allowed; note, though, that adding additional copies of a detector that is already present in the rule does not change its truth values, and thus does not affect the likelihood.) The length of the compound rule is then $n_i = \sum_{j=0}^{n_\mathcal{D}} z_j^{(i)}$.

For implementation purposes, rather than store each vector $z^{(i)}$ explicitly, we represent the rules as lists of detectors objects, which are kept sorted according to an ordering on $\mathcal{D}$. Conceptually, the particular choice of ordering is irrelevant; in practice, we order the detectors in $\mathcal{D}$ lexicographically by their corresponding vectors of truth values.

#### 1.6 Prior probability distribution on rule sets

We place a prior probabiliy distribution $\pi(R)$ on the rule set, which favors parsimonious rules and encodes prior phylogenetic information. The prior probability distribution $\pi(R)$ can be described by the following generative process:

- First, decide whether the rule set is empty. With probability $\Theta_0$, it has length zero; otherwise, it has some nonzero length.
- If the set is nonempty, generate it according to the following procedure:
  1. Set its length to $m$, where the excess length $m-1$ is drawn from a truncated negative binomial distribution with parameters $\alpha_m$ and $\beta_m$ and maximum value $m_{\max} - 1$.
  2. Draw the excess lengths $n_i - 1$ of each rule independently from a truncated negative binomial distribution with parameters $\alpha_n$ and $\beta_n$ and maximum value $n_{\max} - 1$.
  3. Finally, for each rule $\rho_i$, draw $n_i$ detectors from a prior distribution over $\mathcal{D}$ discussed below. (More precisely, draw the vector $z^{(i)}$ from a multinomial distribution with number of trials $n_i$ and event probabilities $p_1, p_2, \ldots, p_{n_\mathcal{D}}$ given below.)

We can write the prior in the sorted tuple format described above as:

$$\pi(R) = \begin{cases} \Theta_0, & m = 0 \\ (1-\Theta_0)\, P_m(m-1) \cdot & \\ \quad \prod_{i=1}^{m} \left(P_n(n_i - 1) M_i p(r_{i1}) p(r_{i2}) \ldots p(r_{in_i})\right), & m > 0 \end{cases} \tag{2}$$

where $p(r_{ij})$ is the probability of detector $r_{ij}$ under the prior discussed below, $M_i$ is the multinomial coefficient $M_i = n_i!/(\prod_{j=0}^{n_\mathcal{D}} z_j^{(i)}!)$, and $P_m$ (respectively $P_n$) is the probability mass function of the truncated negative binomial distribution with minimum value 0, maximum value $m_{\text{max-1}}$ ($n_{\max} - 1$), and parameters $\alpha_m, \beta_m$ ($\alpha_n, \beta_n$).

For ease of exposition later, we define

$$\pi_S(R) = \begin{cases} \Theta_0, & m = 0 \\ (1-\Theta_0)\, P_m(m-1) \prod_{i=1}^{m} \left(P_n(n_i - 1)\right), & m > 0, \end{cases} \tag{3}$$

the contribution to the prior probability of $R$ determined solely by its structure (length and subrule lengths), independent of the particular detectors it contains.

Below we describe each component of the prior and the choice of associated parameters.

**Prior on the probability that the rule set is empty** We assume:

$$\Theta_0 \sim \text{Beta}(a_\Theta, b_\Theta).$$

By choosing $a_\Theta$ and $b_\Theta$ we may adjust the strength of our prior belief that no association exists between the microbiome data and host status. Typically we set $a_\Theta = 0.5$, $b_\Theta = 0.5$, for a 50% prior probability (agnostic) that the list is empty.

**Negative binomial priors for rule lengths and total number of rules** The prior distributions of the number of rules and the number of detectors within each rule are set by default to promote interpretable results via parsimonious rules. Specifically, we take $\alpha_m = 0.5$, $\beta_m = 2.0$, $\alpha_n = 2.0$, and $\beta_n = 4.0$, leading (in the limit of high $m_{\max}$ and $n_{\max}$) to a mean number of rules equal to 1.25 and mean number of detectors per rule equal to 1.5, with 98.7% of rule sets containing 3 or fewer rules, and 97.3% of rules containing 3 or fewer detectors.

The choice of different parameters for the distributions of the number of rules and the number of detectors per rule reflects our belief that a rule set containing one rule with two detectors is generally more straightforward to interpret than a rule set containing two rules, each with one detector. (To support this intuition, we note that in the former case, the rule set divides the population into two groups with different probabilities of the outcome of interest, while in the latter case, four subgroups of the population all have different outcome probabilities, a more complex situation.)

##### 1.6.1 Prior distribution over detectors

For each detector $r^{(k)}$ in $\mathcal{D}$, we assign a prior probability

$$p(r^{(k)}) \propto G(r^{(k)})w(r^{(k)})$$

where $G(r^{(k)})$ depends only on the clade to which the detector applies, and $w(r^{(k)})$ depends only on the duration of the associated time window. These functions are described below. (We assume that all threshold values are equally likely, and that detectors are equally likely to refer to a variable's slope or its average, and equally likely to have 'above' or 'below' direction.)

##### 1.6.2 Phylogenetic prior

We assign to any detector $r^{(k)}$ applying to the aggregated abundance of the descendants of node $i$ a factor

$$G(r^{(k)}) = n(\log L_i; \mu_L, \sigma_L^2)$$

where $n(\cdot; \mu_L, \sigma_L^2)$ is the probability density function of the standard distribution with mean $\mu_L$ and variance $\sigma_L^2$. Here, $L_i$ is the total length of all the edges in the subtree descending from node $i$, plus the length of the edge connecting node $i$ to its parent. The specification in terms of $\log L_i$ is natural, as $\log L_i$ roughly correlates with a node's level in the phylogenetic tree.

Here, our goal is not to specify any preferred phylogenetic level by default, but instead to establish a non-informative or weakly-informative prior, to allow a suitable level (or agnosticism towards phylogenetic level, as appropriate) to be learned from the data. We acheive this as follows. Let $L_l$ and $L_h$ equal the 2.5th and 97.5th percentiles of the *logarithms* of the weights $L_i$ associated with all nodes of the tree and $\Lambda_L$ be the median of the logarithms, and set $\Delta_L = L_h - L_l$. We take

$$\mu_L \sim \mathrm{N}(\Lambda_L, 50\Delta_L^2)$$
$$\sigma_L \sim \mathrm{Uniform}(0, 25\Delta_L).$$

##### 1.6.3 Prior on time window length

We assign to any detector $r^{(k)}$ applying to a time window of length $W_k$ a factor

$$w(r^{(k)}) = B(W_k/((1+\epsilon_w)T_{\text{experiment}}); \alpha_w, \beta_w)$$

where $B(\cdot; \alpha_w, \beta_w)$ is the probability density function of the beta distribution with parameters $\alpha$ and $\beta$. We set

$$\alpha_w = c_w f_w$$
$$\beta_w = c_w(1 - f_w)$$

so that $f_w$ is the prior mean window length, expressed as a fraction of the overall experimental duration, under the notional Beta distribution governing the window lengths $W$. (Note that, because of the discretization, the actual prior mean window length may differ from $f_w$.) The factor of $(1+\epsilon_w)$ prevents the factors assigned to detectors that cover the entire duration of the experiment from becoming infinite when $\alpha_w, \beta_w < 1$; we set $\epsilon_w = 0.01$. For $c_w$ and $f_w$, we introduce a second level of prior distributions with hyperparameters:

$$f_w \sim \mathrm{Uniform}(0, 1)$$
$$c_w \sim \mathrm{Exponential}(\lambda_w)$$

with $\lambda_w = 1/c_{w,\text{typical}} = 1/5$.

#### 1.7 Incorporating additional covariates

##### 1.7.1 Motivation

In many studies, additional information about each host, beyond their microbiome composition, is collected, and we may wish to incorporate these variables into our predictive models, to study or control for their effects on the outcome of interest. For example, we may study the relationship between the microbiome and a disease across two populations (e.g., male and female), with different overall disease incidences; in such a case we would want to know whether any association we observe between the microbiome and the disease is simply a proxy for membership in the higher-incidence population.

##### 1.7.2 Modifying the logistic regression model

We incorporate non-time-dependent host covariates into the logistic regression model 1 as follows.

Let $C$ be a matrix of host covariates, such that $C_{ij}$ is the value of variable $j$ for host $i$, $j = 1, \ldots, n_c$.

We first consider the common situation of categorical covariates, encoded as integers. For each such covariate $j$ we specify a default value $\hat{c}_j$. Because the encoding is arbitrary, we can assume without loss of generality that $\hat{c}_j = 0$ and the observed values of variable $j$ run from 0 to $m_j$.

For each $j$, we define a new matrix $\mathcal{C}^{(j)}$ such that, for $1 \leq k \leq m_j$, $\mathcal{C}^{(j)}_{ik} = 1$ if $C_{ij} = k$ and otherwise $\mathcal{C}^{(j)}_{ik} = 0$. Then let

$$\mathcal{C} = \left( C^{(1)} | \ldots | C^{(n_c)} \right).$$

For example, if we have three subjects and two additional variables, with

$$C = \begin{pmatrix} 0 & 1 \\ 1 & 0 \\ 0 & 2 \end{pmatrix}$$

we obtain

$$\mathcal{C} = \begin{pmatrix} 0 & 1 & 0 \\ 1 & 0 & 0 \\ 0 & 0 & 1 \end{pmatrix}.$$

Now, in 1, we replace $A(R, x)$ with

$$A(R, C, x) = (A(R, x)|\mathcal{C}).$$

Effectively, for each variable $j$, we have augmented $A(R)$ with a incomplete one-hot encoding of $j$ that leaves out the column that would correspond to the 'default' value $\hat{c}_j$. This exclusion is important, because otherwise severe collinearity issues would arise.

To extend this approach to include discrete or continuous quantitative variables, we carry out the same process, but where column $j$ of $C$ includes such a variable, we simply set $\mathcal{C}^{(j)}$ equal to column $j$ of $C$, effectively inserting each such column of covariates directly into $A(R, C)$.

Note that we will now have regression coefficients $\beta$ corresponding to each rule in our rule set $R$, a coefficient associated with the constant/intercept term, and a coefficient for each quantitative host covariate and each non-default value of each categorical host covariate observed anywhere in our population; the latter coefficients indicate the change in disease risk associated with those values (relative to the default value) conditional on $R$.

Importantly, host covariates are always present in the regression calculation, but the prior distribution over rule sets penalizes the inclusion of rules which do not improve the likelihood; thus, under the MITRE posterior distribution rules which do not provide additional information about the outcome of interest beyond that encoded in the host covariates will be unlikely to be included in the rule list.

### 2 Inference algorithm

#### 2.1 Outline

Our objective is to infer the posterior distribution of rule sets $R$ and regression coefficients $\beta$ given the data. To accomplish this, we use a custom Markov Chain Monte Carlo (MCMC) algorithm with the following iterated steps:

1. Update the coefficients $\boldsymbol{\beta}$, drawing new values from their conditional distributions using a data augmentation technique.

2. Update the rule set $R$ via a Metropolis-Hastings step which proposes one of the following modifications to the structure of $R$:

    - Replacing one detector with another detector from $\mathcal{D}$
    - Moving a detector from one rule to another
    - Adding a detector to the set
    - Removing a detector from the set.

3. Update the hyperparameters controlling the prior distribution $\pi(R)$ using Metropolis-Hastings steps.

For detailed discussion of each of these steps, see the Appendix.

##### 2.2 Assessment of mixing

We confirmed that the MCMC algorithm achieved adequate mixing by inspecting plots of the likelihood, prior probability, rule set length (total number of detectors), auxiliary variables $\boldsymbol{\omega}$, and parameters $f_w$, $c_w$, $\mu_L$, and $\sigma_L$ versus MCMC iteration; and by calculating for those parameters the convergence metric[2] $\hat{R}$ using multiple Markov chains for one representative calculation with each data set, confirming that the values were consistent with convergence ($\lessapprox 1.1$) after the standard number of iterations used for calculations with the dataset in question (25000-100000; see main text).

These plots are generated automatically for each run by setting the `mixing_diagnostics` option in the `postprocessing` section of the MITRE configuration file to True; calculation of the $\hat{R}$ values from multiple Markov chains is discussed in the MITRE software manual.

#### 3 Preparing input data for MITRE

##### 3.1 Introduction: MITRE accepts many types of input data

The MITRE model is flexible, and may be applied to microbiome abundance data in a variety of forms. For instance, data could be counts data for OTUs identified based on 16S amplicon sequencing; relative abundances predicted by marker gene profiling of shotgun metagenomics data; absolute abundances determined by qPCR, etc.

Generally, the only constraint is that data should be on, or easily convertible to, a scale where it is sensible to define the abundance of a clade as the sum of the abundances of its descendants. Even this restriction may be avoided if the user provides data for higher clades manually rather than invoking MITRE's automatic aggregation features.

[2] Gelman et al., *Bayesian Data Analysis* 3rd Ed, CRC Press, 2013, p. 284-285.

In sections 3.2, 3.3, and 3.4, as an illustration of MITRE's capabilities, we outline the recommended preprocessing, filtering, and annotation workflow for the common case of 16S sequencing counts data, with a phylogenetic tree defined by placing OTUs on a reference tree using `pplacer`. Note that most filters and transformations described in section 3.2 may be readily applied to other data types as well. In section 3.5 we discuss some special aspects of MITRE's processing of taxonomic profiles generated by Metaphlan.

#### 3.2 16S data processing pipeline

The following procedure for filtering and processing OTU counts data before supplying it as input to the MITRE algorithm is recommended. (The description of this process and recommended parameter values given in the main text of the paper is reproduced here in the interest of providing a self-contained description of the MITRE approach in this document.) Note that each described step can be carried out automatically by the MITRE software before beginning the inference process, if it is indicated in the configuration file.

##### 3.2.1 Filtering steps before aggregation

1. To remove potentially spurious OTUs, we discard data for all OTUs with less than $N_{\text{counts,OTU}}$ counts observed across all samples (recommended value: 10.)

2. To exclude samples where coverage is so low that abundance estimates for uncommon OTUs may be unreliable, discard data from all samples with less than $N_{\text{counts,sample}}$ total counts observed across all remaining OTUs (recommended value: 5000 for HiSeq/MiSeq data.)

3. If desired, to analyze only a particular time period (because, e.g., samples are not available outside that period for the majority of subjects,) discard all samples before time $t_i$ and after time $t_f$ (by default, $t_i$ is the time of the earliest available sample, and $t_f$ the time of the latest sample.)

4. To exclude subjects for whom microbiome dynamics cannot be resolved at the desired temporal resolution throughout the entire study, discard subjects with too few, or too sparse, observation points, by dividing

the duration of the study into $N_{\mathrm{w,filter}}$ equal pieces and keeping only subjects with at least $N_s$ samples in any $N_c$ consecutive such pieces. Default values are $N_{\mathrm{w,filter}} = 10$, $N_s = 2$, and $N_c = 1$; note that for data with very inconsistent sampling, these parameters must be chosen judiciously to maximize the number of subjects included while allowing an acceptable level of temporal resolution.

###### 3.2.2 Phylogenetic aggregation

Next, counts data are converted to relative abundance data for each sample, and, for each node in the phylogenetic tree, a relative abundance estimate is obtained by summing the relative abundances of its children. (Note that the standard approach is to determine abundances relative to the total bacterial community, but by setting an appropriate configuration option, abundances may be calculated relative to the abundance of a specified taxonomic group.)

###### 3.2.3 Filtering steps after aggregation

The following filtering steps are then applied to all clades (both OTUs and higher nodes in the tree):

1. To exclude low abundance clades (for which abundance estimates may be inaccurate) or infrequently observed clades (which we expect are unlikely to be explanatory, though higher clades including them may be), discard all clades except those which exceed a threshold abundance a in at least $N_a$ consecutive samples in at least $N_i$ subjects. Typically, $N_a = 2$. Appropriate values for $a$ and $N_i$ depend on the number of subjects and typical reads per sample; for studies with 10-150 subjects and average reads per sample on the order of $10^4$, we recommend $a = 10^{-4}$ and $N_i = 4$.

2. To minimize redundancy, discard all clades corresponding to nodes in the phylogenetic tree with exactly one remaining child clade, as their dynamics are often very similar to those of their children.

Note that, when a large number of taxa are considered, the detector pool becomes large and the computational cost of the inference algorithm grows; if necessary, it is recommended to increase the stringency of these steps to keep the total number of taxa below 500.

#### 3.3 Generating phylogenetic trees from pplacer output

##### 3.3.1 Introduction

The MITRE model outlined above is agnostic as to how the phylogenetic tree relating the OTUs is obtained. The software implementing the model currently assumes that such trees will be obtained by placing representative sequences from each OTU on a reference tree of known 16S sequences using the `pplacer` package[3] in maximum likelihood mode. Here, we describe how we processs the `pplacer` output to obtain one tree relating the OTUs.

##### 3.3.2 The pplacer output format

Only one output file from `pplacer` need be supplied to MITRE: the `.jplace` file, which contains, in JSON format, a representation of the reference tree onto which sequences have been placed, and information about one or more inferred possible placements for each sequence, including among other data:

- The edge on the reference tree to which the query sequence might be placed
- The 'pendant length' of the edge that would connect the sequence to its point of attachment to the reference tree
- A 'likelihood weight ratio' between 0 and 1, indicating the relative confidence of this placement compared to alternatives.

Although the likelihood weight ratios are not true Bayesian posterior probabilities we interpret them in an analogous fashion. Note in particular that the likelihood weight ratios for the possible placements of a particular query sum to 1.0.

##### 3.3.3 Obtaining a single tree

While the `pplacer` output includes, in general, multiple possible positions for each query sequence, MITRE requires a single tree relating the OTUs. We obtain such a tree by placing each query sequence as a child of the

[3] Matsen, F. A., Kodner, R. B. & Armbrust, E. V. "pplacer: linear time maximum-likelihood and Bayesian phylogenetic placement of sequences onto a fixed reference tree". *BMC Bioinformatics* 11, 538 (2010).

'lowest' edge such that we are confident that the sequence should be placed somewhere on the subtree descending from that edge, proceeding for each query sequence as follows:

- To each edge on the reference tree, assign a score equal to the sum of the likelihood weight ratios of all candidate placements of the query to that edge, or any edge descending from that edge.
- Find the edge with the lowest score that is still greater than 0.5. We will attach the query to this edge, with a pendant length equal to the average of the pendant lengths of the candidate placements, weighted by their likelihood weight ratios.
- We do not know at what position along the edge the query should attach, but for our purposes it is sufficient to attach the query as a child node of the node at the lower end of the edge.

After carrying out this process for all query sequences, we prune the tree, removing nodes and edges with no query sequences as descendants, and then collapsing internal nodes with only one child node. After this pruning process we calculate the subtree lengths $L_i$ used in the phylogenetic prior (section 1.6.2.) (Note that these weights are not subsequently recalculated when one clade or another is excluded in the filtering process of section 3.2.)

#### 3.4 Taxonomic annotations

Although it is not required for the operation of the software, labeling each OTU or higher clade according to a taxonomic classification facilitates interpretation of the results. Optionally, given appropriate input files, MITRE can perform such a labeling; it then writes a table of labels for each OTU/clade and displays the labels as appropriate in the interactive graphical interface. Three methods of label generation are possible:

**pplacer-based** In this method, additional files from the `pplacer` reference package used to perform phylogenetic placement are supplied to MITRE. Nodes in the reference tree are labeled according to the lowest common taxonomic classification of the reference species which descend from them; then OTUs are labeled according to the classication of their parent nodes.

**Table-based** In this method, MITRE is supplied a table of predicted classifications for each OTU sequence, as can be generated with `mothur`'s RDP-based classification tool, or `dada2`'s RDP-based classification plus exact sequence matching capability. In this case, OTUs are labeled according to the table; then higher nodes in the tree are labeled according to the lowest common predicted classification of the OTUs which descend from them.

**Hybrid** In practice, the `pplacer`-based method gives more informative descriptions of higher nodes, while the table-based method often (but not always) gives more specific descriptions of OTUs. The hybrid method uses all the input files from the other two methods to first label the nodes and OTUs following the `pplacer`-based method, then updates OTU labels from the table for those OTUs for which a species-level assignment is available, offering the advantages of both approaches.

#### 3.5 Notes on working with Metaphlan results

Metaphlan output files provide relative abundance estimates (on a percentage scale) for taxonomic groups ranging from below the species level to kingdoms. As such it is not necessary for MITRE to explicitly carry out phylogenetic aggregation. Filtering taxa based on abundance, as described for 16S data, is still recommended to ensure that the total number of variables remains manageable.

MITRE will automatically infer a tree structure that relates the taxa appearing in a Metaphlan output file, but it is not possible to automatically determine biologically meaningful edge lengths for this tree. Thus, by default, the weights $L_i$ of section 1.6.1 are set equal to 1.0 for all nodes in the tree; alternatively, by setting a configuration option it is possible to set all edge lengths to 1.0 and then determine the weights $L_i$ as usual.

### 4 Data processing settings used for particular datasets

#### 4.1 Vatanen et al.

For Vatanen et al., demographic and outcome data for each subject and FASTQ files of 16S rRNA amplicon sequences (175bp paired-end sequences of the V4 region generated on the Illumina HiSeq 2500 platform, average depth 57000 across 1584 total samples) were downloaded from the DIABIMMUNE project web site[4] and processed as described above. Preprocessing settings were $N_{\text{counts,otu}} = 10$, $N_{\text{counts,sample}} = 5000$ (the defaults), $t_i = 27$ days (the date of the earliest sample), $t_f = 900$ days (because, within the Russian cohort in particular, only a small fraction of subjects had any later samples available), $N_{\text{w,filter}} = 9$, $N_s = 2$, $N_c = 4$, $a = 10^{-4}$, $N_a = 2$, and $N_i = 11$. The choices of $N_{\text{w,filter}}$, $N_s$, and $N_c$ allowed approximately the maximum possible temporal resolution consistent with including more than 50% of subjects in the analysis (in total, 111 subjects were retained after filtering). The MITRE detector pool was generated using $N_w = 9$, $W_{\text{min}}$=0 days, and $W_{\text{max}} = 450$ days (half the duration of the study).

#### 4.2 Bokulich et al.

For Bokulich et al., FASTQ files of 16S rRNA amplicon sequences (150 bp sequences of the V4 region generated on the Illumina MiSeq platform, average depth 21360 across 762 total samples) were downloaded from ENA and metadata from QIITA (`qiita.ucsd.edu`, study 10249) and processed as described above. MITRE preprocessing settings were $N_{\text{counts,otu}} = 10$, $N_{\text{counts,sample}} = 5000$ (the defaults), $t_i = 0$ and $t_f = 375$ days (covering the portion of the study in which samples were collected monthly), $N_{\text{w,filter}} = 12$, $N_s = 1$, $N_c = 2$, $a = 10^{-4}$, $N_a = 3$, and $N_i = 10$. The choices of $N_{\text{w,filter}}$, $N_s$, and $N_c$ allowed approximately the maximum possible temporal resolution consistent with including in the analysis more than 90% of subjects for whom samples were available through the entire first year of life (note that the desired fraction of subjects retained is greater here, and below, than for Vatanen et al, because the total number of subjects is smaller). 35 subjects were retained after filtering. The MITRE detector pool was generated using

[4] `https://pubs.broadinstitute.org/diabimmune/three-country-cohort/`

$N_w = 12$, $W_{\min} = 0$ days, and $W_{\max} = 180$ days (half the duration of the study.)

#### 4.3 Di Giulio et al.

For Di Giulio et al., FASTQ files of 16S rRNA amplicon sequences (V3-V5 region sequenced on the 454 FLX platform) were obtained from the authors. Sequences from vaginal swab samples taken during pregnancy (767 total samples, average depth 5600) were processed as described above. MITRE preprocessing settings were $N_{\text{counts,otu}} = 10$, $N_{\text{counts,sample}} = 1500$, $t_i = 140$ and $t_f = 210$ days, $N_{\text{w,filter}} = 1$, $N_s = 1$, $N_c = 1$, $a = 0.005$, $N_a = 2$, and $N_i = 4$; the MITRE detector pool was generated using $N_w = 1$, $W_{\min} = 0$ days, and $W_{\max} = 100$ days (half the duration of the study.) Non-default values of $N_{\text{counts,otu}}$ and $a$ reflected the lower average sequencing depth in this 454-based study. 36 subjects were retained after filtering. Dates of the first and last available samples varied widely across patients in this study; the use of a single time window between 20 and 30 weeks gestational age maximized the number of patients in the premature delivery group included in the analysis.

#### 4.4 David et al.

For David et al., FASTA files of 16S amplicon sequences (100bp paired-end sequences of the V4 region generated on the Illumina HiSeq platform; average depth 43600 across 235 total samples) were downloaded via MG-RAST and processed using mothur as described above. Because these files contain concatenated non-overlapping forward and reverse reads, the Gotoh alignment method with no gap extension penalty was used for alignment against the reference database. MITRE preprocessing settings were $N_{\text{counts,otu}} = 10$ and $N_{\text{counts,sample}} = 5000$ (the defaults), no filtering of subjects based on sampling frequency, and the default values $a = 0.001$, $N_a = 2$, and $N_i = 4$. Processing with mothur resulted in a large number of OTUs so the larger value of $a$ was chosen to maintain tractability. The MITRE detector pool was generated using $N_w = 10$, $W_{\min} = 0$ days, and $W_{\max} = 16$ days (the duration of the study.) Although the same subjects participated in the plant and animal arms of the study, we analyzed them as though they were independent populations.

##### 4.5 Kostic et al.

For Kostic et al, metaphlan results and WGS sample metadata from was downloaded from the DIABIMMUNE website (`https://pubs.broadinstitute.org/diabimmune/t1d-cohort`) and minimally reformatted using a script distributed with MITRE. MITRE preprocessing settings were $t_i = 0$ and $t_f = 1000$ days, $N_{\text{w,filter}} = 5$, $N_s = 1$, $N_c = 2$, $a = 0.001$, $N_a = 3$, and $N_i = 10$; the MITRE detector pool was generated using $N_w = 5$, $W_{\min} = 1$ days, and $W_{\max} = 500$ days (half the duration of the study.)

### 5 Postprocessing: generating summaries and visualizations from MITRE posterior samples

##### 5.1 Introduction

Each individual MITRE rule set is designed to be readily interpretable. The inference algorithm described above generates collections of typically tens of thousands of samples from the posterior distribution over the space of possible rule sets, however, which are less amenable to direct inspection. To interpret the posterior samples, MITRE offers four tools: a single-rule-set point estimate; a clustering of the samples, ranked by posterior probability; Bayes factors quantifying the evidence for particular hypotheses about the true rule set; and an interactive visualization tool.

##### 5.2 Point estimate

The point estimate is a single rule set $R^*$ with associated coefficients $\boldsymbol{\beta}^*$ that is chosen to be a high-likelihood (i.e., parsimonious yet accurately predictive) rule set that is broadly representative of the posterior samples.

It is determined according to the following procedure:[5]

- If the posterior probability that $R$ is empty is greater than 0.5, the point estimate is the empty rule set.

[5] This point estimation procedure is similar to and inspired by the estimation procedure applied to a different model by Letham et al, “Interpretable classifiers using rules and Bayesian analysis: Building a better stroke prediction model.” *Ann. Appl. Stat.* 9, 13501371 (2015).

- Otherwise, to obtain a representative non-empty summary list, we determine the posterior mode $d^*$ of the total number of detectors in $R$, and take $R^*$ to be the rule set with the highest posterior probability among all sampled rule sets which contain $d^*$ total detectors. The point estimate coefficients $\boldsymbol{\beta}^*$ are the highest posterior probability sampled coefficients associated with $R^*$.

#### 5.3 Clustering

##### 5.3.1 Outline

To summarize the diversity of alternative rule sets in the posterior samples, MITRE clusters together similar rule sets and ranks the clusters by posterior probability. Broadly, we consider two rule sets 'similar' if they contain rules made up of detectors that apply to the same function (slope or average) of the same variables, in the same or significantly overlapping time windows. In detail, we take a bottom-up approach, first clustering detectors, then using that clustering to establish a clustering of rules, then finally establishing a cluster of rule sets. The specific procedure is described below.

##### 5.3.2 Clustering detectors

We sort all detectors in the posterior samples by the number of times they appear. The most common detector is assigned to the first cluster. We proceed through the detectors which have not yet been added to any cluster in descending order of frequency and decide whether to add each one to the cluster, doing so if both of the following conditions are met:

- The candidate detector applies to the same function (slope/average) of the same variable, and has the same direction (above/below), as all the detectors added to the cluster so far.

- Comparing the time window of the candidate detector to the time window of each detector already in the cluster, we find in each case that at least 50% of the shorter time window is contained within the longer window.

We then form a new cluster with the most-frequent detector not yet added to a cluster, and repeat the process until all detectors have been assigned to a cluster.

##### 5.3.3 Clustering rules

We consider two rules to be equivalent if either of the following conditions is met:

- Their representations in terms of the clusters to which their component detectors belong (e.g., two detectors from cluster 1 and one detector from cluster 18) are identical;
- Each rule contains only one detector, and the *opposite* of one of the detectors (formed by switching the direction from 'above' to 'below' or vice versa) belongs to the same cluster as the other.

This equivalence relation naturally partitions the set of observed rules into equivalence classes, which we take as our clustering of the rules.

##### 5.3.4 Clustering rule sets

Similarly, we consider two rule sets to be equivalent if they contain the same numbers of rules from the same rule clusters (e.g., both contain a rule from cluster 1, a rule from cluster 2, and two rules from cluster 5.) The equivalence classes under this relation form the clustering of rule sets.

#### 5.4 Bayes factors

##### 5.4.1 Introduction

In addition to identifying one or more high-posterior-probability rule sets, investigators may wish to ask questions such as:

- How strong is the evidence that the true rule set is nonempty?
- How strong is the evidence that the true rule set involves a particular clade or OTU?
- How strong is the evidence that a particular OTU or some ancestral clade, at a particular time, is associated with the outcome of interest?

MITRE answers such questions by calculating *Bayes factors*, which offer a formal method for quantifying the strength of the evidence for one model over an alternative in a Bayesian context.[6]

[6] Kass, R. E. & Raftery, A. E. "Bayes Factors". *J. Am. Stat. Assoc.* 90, 773795 (1995).

We can calculate a Bayes factor for any proposition which can be put in the form of a hypothesis that the true rule set $R$ belongs to a particular subset of the space of all possible rule sets: for example,

- the space of nonempty rule sets,
- the space of rule sets including at least one detector that applies to a particular clade or OTU,
- the space of rule sets including at least one detector that applies to a particular OTU or any of its ancestors, and has a time window overlapping a time period of interest.

Specifically, if our hypothesis is that $R$ belongs to the set of rule sets $S$, the Bayes factor in favor of this hypothesis is

$$\frac{P(R \in S|\mathbf{y})}{P(R \notin S|\mathbf{y})}\frac{P(R \notin S)}{P(R \in S)},$$

that is, the ratio of the posterior odds of the hypothesis to the prior odds. One conventional interpretation of such factors follows a logarithmic scale, with values greater than 10 indicating strong evidence in favor of the hypothesis.[7]

Below, we discuss the procedure through which MITRE calculates Bayes factors for various hypotheses, including those discussed above, from the posterior samples generated by the inference procedure.

##### 5.4.2 For the empty rule

The prior probability that $R$ is empty is simply $a_\Theta/(a_\Theta + b_\Theta)$; thus, if the fraction of posterior rule set samples which are empty is $f$, the estimated Bayes factor for the empty rule set is

$$\frac{f}{1-f}\frac{a_\Theta + b_\Theta}{a_\Theta}.$$

(and the Bayes factor for the hypothesis that the rule set is non-empty is the reciprocal of this quantity.)

This factor gauges the strength of the evidence for an association between the microbiome dynamics and host status; if it is not substantially less than 1, caution is warranted in interpreting other results.

[7] Ibid.

##### 5.4.3 For individual detectors

The strength of the evidence that any individual detector $r^*$ is involved in the true rule set $R$ is rarely of key interest on its own, but the procedure for evaluating such a Bayes factor forms the basis for evaluating Bayes factors for more complex and interesting hypotheses.

First, we estimate the prior probability $p(r^*)$ associated with the detector, defined in section 1.6.1. (Recall that conceptually, this is the probability that a randomly chosen detector in a rule set drawn from the prior distribution over rule sets happens to be $r^*$.) This is done by drawing repeated samples of the hyperparameters $\mu_L$ and $\sigma_L$ of the phylogenetic prior (section 1.6.2) and $c_w$ and $f_w$ of the time window prior (section 1.6.3) from their respective higher-level prior distributions and evaluating $p(r^*)$ under each sampled set of hyperparameters, then averaging the values to obtain an estimate $\hat{p}(r^*)$ of the prior probability associated with the detector, marginalizing out those hyperparameters. (By default, MITRE uses 10000 samples of the hyperparameters to make this estimate; by reusing these samples we may relatively efficiently obtain estimated prior probabilities for each detector in $\mathcal{R}$ if desired.)

Next, we estimate the prior distribution of the total number of detectors in $R$. Note that the maximum number of detectors is $r_{\max} = m_{\max} n_{\max}$ in the notation of section 1.6. We estimate the probability of each possible number of detectors, conditional on the rule set being nonempty, by repeatedly drawing numbers of rules $m$ and numbers of detectors in each rule from their prior distributions, counting the number of times a rule set with each possible total number of detectors is drawn, and dividing those counts by the number of samples. (By default MITRE uses 100000 sampled rule sets for this estimation procedure).

Weighting these probabilities by the prior probability $b_\Theta/(a_\Theta + b_\Theta)$ that the rule set is non-empty, we obtain estimates $\hat{p}_d(1), \hat{p}_d(2), \ldots, \hat{p}_d(r_{\max})$ of the overall prior probability of each possible number of detectors. Let $p_d(0) = \hat{p}_d(0) = a_\Theta/(a_\Theta + b_\Theta)$ be the prior probability that the rule set is empty.

The prior probability that a rule set with $d$ total detectors does *not* contain any copies of $r^*$ is evidently

$$(1 - p(r^*))^d$$

for $d \geq 0$. It follows that the marginal prior probability that a rule set *does* contain at least one copy of $r^*$ is

$$P(r^* \in R) = 1 - \sum_{i=0}^{r_{\max}} p_d(i)(1 - p(r^*))^i \approx 1 - \sum_{i=0}^{r_{\max}} \hat{p}_d(i)(1 - \hat{p}(r^*))^i. \quad (4)$$

It is straightforward to tabulate the fraction $f^*$ of the posterior rule set samples that contain at least one copy of $r^*$, leading to the estimated Bayes factor

$$B_{r^*} = \frac{f^*}{1 - f^*} \frac{1 - P(r^* \in R)}{P(r^* \in R)} \quad (5)$$

for the hypothesis that the true rule set contains at least one copy of $r^*$.

##### 5.4.4 For higher-level groupings of detectors

If instead we are interested in the evidence that the true rule set contains at least one detector belonging to a defined set $S \subset \mathcal{R}$, we can estimate the Bayes factor for that hypothesis using a similar procedure.

First, for each $r \in S$, calculate an estimated prior probability $\hat{p}(r)$ as above. Then

$$\hat{p}(S) = \sum_{r \in S} \hat{p}(r)$$

is the estimated prior probability that any given detector in a rule set drawn from the prior distribution over rule sets belongs to $S$.

Defining and evaluating as above the estimated prior probabilities

$$\hat{p}_d(0), \hat{p}_d(1), \hat{p}_d(2), \ldots, \hat{p}_d(r_{\max})$$

that the rule set will contain $0, 1, \ldots, r_{\max}$ total detectors, the same argument as above shows that the marginal prior probability that $R$ contains at least one detector belonging to $S$ is

$$P(R \cup S \neq \emptyset) \approx 1 - \sum_{i=0}^{r_{\max}} \hat{p}_d(i)(1 - \hat{p}(S))^i. \quad (6)$$

If $f_S$ is the fraction of posterior rule set samples containing at least one detector belonging to $S$, we obtain the estimated Bayes factor

$$B_S = \frac{f_S}{1 - f_S} \frac{1 - P(R \cup S \neq \emptyset)}{P(R \cup S \neq \emptyset)} \quad (7)$$

for the hypothesis that the true rule contains at least one copy of $r^*$.

##### 5.4.5 A note on the accuracy of Bayes factor estimates

The estimated Bayes factors derived above rely on approximations of both the prior probabilities of each hypothesis and the posterior probabilities, the latter obtained by counting the fraction of a collection of rule sets, sampled using the MITRE inference procedure, that satisfy a particular hypothesis. Clearly, this procedure will never give a nonzero posterior probability estimate less than $1/n_{\text{samples}}$, the reciprocal of the number of samples available, and will in general not lead to accurate estimates of posterior probabilities $\lessapprox 1/n_{\text{samples}}$. We recommend using 10000-50000 samples for practical MITRE calculations. However the total number of detectors $n_{\mathcal{R}}$ is often over 100000, and the prior probabilities $p(r)$ associated with each detector will typically be of order $1/n_{\mathcal{R}}$.

Estimated Bayes factors for the hypothesis that relatively unlikely detectors (or sets of detectors) are represented in the true rule set thus may be inaccurate. In particular, the estimated posterior odds that the rule set contains a particular detector $r*$ that appears only in a single sample will typically be greater than the prior odds that that detector appears in the rule set, leading to a potentially inflated Bayes factor estimate.

Generally this does not lead to practical difficulties, for two related reasons. First, we usually are most interested in hypotheses with higher posterior probabilities, for which estimates will be more accurate. Second, we primarily use Bayes factors to rank and compare the evidence that variables or variable/time-window combinations are invovled in the true rule set, and thus are most concerned with the relative, not absolute, values of the estimates.

However, users who wish to obtain accurate Bayes factors for low-posterior-probability hypotheses will wish to increase $n_{\text{samples}}$ beyond the usually recommended values.

#### 5.5 Interactive visualization

The MITRE interactive visualization allows the user to quickly identify time periods and clades associated with a particular host status. Here we discuss a few of its key features and describe in detail the calculations performed to support them.

The primary element of the visualization tool is a grid in which each each column corresponds to one of the $N_w$ time windows defined in section 1.4

and each row corresponds to an OTU (or a low-level clade whose child OTUs have been removed by filtering). The OTUs are arranged according to the phylogenetic tree relating them, which is drawn next to the grid.

Each cell in the grid is colored, heat map style, according to the estimated Bayes factor for the hypothesis that the true rule set $R$ contains at least one detector which:

- applies to the variable corresponding to the row, or any clade containing it, and
- applies to a time window overlapping the time window represented by the column.

Call the set of detectors meeting these conditions $S_{\text{cell}}$.

The user may click on a cell to see a list of the detectors in $S_{\text{cell}}$, ranked by the estimated Bayes factors for each individual detector (as in section 5.4.3.)

Selecting a detector from this list generates plots of the data for the relevant variable for subjects with positive and negative outcomes, with the time window and threshold value of the detector superimposed. If available, the taxonomic annotation of the relevant variable is also displayed.

All relevant Bayes factor estimates are pre-tabulated by MITRE when the standalone HTML file for the visualization is generated.

### 6 An example rule set with interactions inferred from real data

Vatanen et al. (2015) tracked the gut microbiome composition of Russian, Estonian and Finnish infants through the first few years of life. When MITRE was applied to classify subjects in the study according to whether or not they were members of the Russian cohort (the same classification task used in the parameter sensitivity discussion in the Appendix,) the point estimate rule set MITRE learned was: "If, between day 27 and day 415, the average abundance of group 14676 (in the family Enterococcaceae) is below 2.7%, and between day 124 and day 415, the average abundance of group 13204 (in the order Clostridiales) is below 2.0%, and between day 512 and day 803, the abundance of OTU 163 (*Eubacterium hallii*) is below 0.55%, the probability that the subject belongs to the Russian cohort is $3 \cdot 10^{-5}$; otherwise it is 0.997." This rule set provides an illustrative example of MITREs ability to achieve high predictive accuracy while maintaining interpretability: it expresses a nonlinear interaction across three different variables and time windows, but can be easily understood by the user, and achieves an accuracy for this classification task comparable to that of an uninterpretable random forest model that aggregates the predictions of over 1000 decision trees (figure 1(c).)
