## Supplemental Details for "MITRE: predicting host status from microbiota time-series data"

**3 Parameter sensitivity assessment** **18**

### Chapter 1

### Technical description of the inference algorithm

As discussed in the supplementary note, MITRE inference employs a Markov Chain Monte Carlo (MCMC) algorithm with the following iterated steps (using the notation introduced in chapter 1 of the supplementary note):

1. Update the coefficients $\boldsymbol{\beta}$, drawing new values from their conditional distributions using a data augmentation technique.

2. Update the rule set $R$ via a Metropolis-Hastings step which proposes one of the following modifications to the structure of $R$:

   - Replacing one detector with another detector from $\mathcal{D}$
   - Moving a detector from one rule to another
   - Adding a detector to the set
   - Removing a detector from the set.

3. Update the hyperparameters controlling the prior distribution $\pi(R)$ using Metropolis-Hastings steps.

In this chapter we discuss each step in detail. Some derivations are deferred to chapter 2.

#### 1.1 Sampling the logistic regression coefficients

To facilitate inference of the logistic regression coefficients $\boldsymbol{\beta}$, we adopt the data-augmentation approach of Polson, Scott, and Windle[1] and introduce auxiliary variables $\omega_i$, $i = 1, \ldots, n_{\text{subjects}}$, such that, conditional on $\boldsymbol{\beta}$ (and $A$, though we elide the conditioning on $A$ throughout this section for readability), the $\omega_i$ are independent of each other and host status $y_i$, with

$$\omega_i \sim \text{PG}(1, a_i^T \boldsymbol{\beta}) \tag{1.1}$$

where $a_i = (A_{i1}, A_{i2}, \ldots, A_{i(m+1)})$ is row $i$ of $A$ and $\text{PG}(b, c)$ denotes the Polya-Gamma distribution with parameters $b$ and $c$, defined by Polson et al.

Denote the density of $\boldsymbol{\omega}$ under (1.1) as $f(\boldsymbol{\omega}|\boldsymbol{\beta})$. The augmented posterior distribution

$$\pi(\boldsymbol{\beta}, \boldsymbol{\omega}|\mathbf{y}) = \frac{1}{c(\mathbf{y})} \pi(\boldsymbol{\beta}) f(\boldsymbol{\omega}|\boldsymbol{\beta}) \prod_{i=1}^{n_{\text{subjects}}} \Pr(y_i|\boldsymbol{\beta})$$

yields the same posterior density $\pi(\boldsymbol{\beta}|\mathbf{y})$ as the original logistic regression model after marginalizing out $\omega$. The advantage of this formulation is that both of the conditional distributions $\pi(\boldsymbol{\omega}|\mathbf{y}, \boldsymbol{\beta})$ and $\pi(\boldsymbol{\beta}|\mathbf{y}, \boldsymbol{\omega})$ are easy to sample from. The former is simply $f(\boldsymbol{\omega}|\boldsymbol{\beta})$, a combination of independent PG distributions, while the latter can be shown (using the properties of the Polya-Gamma distribution) to be multivariate normal, as follows:

- Define

$$\begin{aligned} \Omega &= \text{diag}(\boldsymbol{\omega}) \\ \boldsymbol{\kappa} &= \left( y_1 - \frac{1}{2}, \ldots, y_{n_{\text{subjects}}} - \frac{1}{2} \right) \\ \mathbf{z} &= \left( \kappa_1/\omega_1, \ldots, \kappa_{n_{\text{subjects}}}/\omega_{n_{\text{subjects}}} \right). \end{aligned}$$

  The likelihood of the observations $\mathbf{y}$ conditioned on $\boldsymbol{\beta}$ and $\boldsymbol{\omega}$ is proportional to

$$\exp\left\{ -\frac{1}{2} (\mathbf{z} - A\boldsymbol{\beta})^T \Omega (\mathbf{z} - A\boldsymbol{\beta}) \right\}.$$

[1] "Bayesian Inference for Logistic Models Using Polya-Gamma Latent Variables", *JASA* 108:1339-1349, 2013. The presentation of the method given here follows Choi and Hobert, "The Polya-Gamma Gibbs sampler for Bayesian logistic regression is uniformly ergodic", *Electron. J. Stat* 7:2054-2064(2013).

We thus have a standard Bayesian regression problem with response $\mathbf{z}$, design matrix $A$, covariance matrix $\Omega^{-1}$, and coefficients $\boldsymbol{\beta}$.

- Proceeding as in the usual case of Bayesian regression with a normal prior on the coefficients, it follows that, in the general case where $\boldsymbol{\beta} \sim N(\mathbf{b}, B)$,

$$\begin{aligned}
\boldsymbol{\beta}|\mathbf{y}, \boldsymbol{\omega} &\sim \mathrm{N}(m_\omega, V_\omega) \\
V_\omega &= \left(A^T \Omega A + B^{-1}\right)^{-1} \\
m_\omega &= V_\omega \left(A^T \boldsymbol{\kappa} + B^{-1}\mathbf{b}\right)
\end{aligned} \tag{1.2}$$

  Here, we have in particular $\boldsymbol{\beta} \sim N(\mathbf{0}, \sigma_\beta^2 I)$, and so

$$\begin{aligned}
V_\omega &= \left(A^T \Omega A + \frac{1}{\sigma_\beta^2} I\right)^{-1} \\
m_\omega &= V_\omega A^T \boldsymbol{\kappa}.
\end{aligned}$$

Using these relationships, we update $\boldsymbol{\beta}$ and $\boldsymbol{\omega}$ in turn 10 times between each update to the rule set structure.

#### 1.2 Sampling rule sets

By proposing to add or remove detectors, swap detectors between rules, or replace a detector in the rule set with some other detector in $\mathcal{D}$, the inference algorithm can potentially explore the entire space of possible rule sets $R$.

The most obvious approach to allow addition would be to implement a very simple Metropolis-Hastings steps which, given a current rule set $R$, adds a detector chosen at random from the pool $\mathcal{R}$ to some position in $R$, forming a new rule set $R'$. However, for any given $R$, we find that a very large fraction of the proposals which could be generated in this way are substantially less probable under the posterior distribution than $R$. (As a rule, the pool $\mathcal{R}$ is very large and most detectors in the pool have no association with the outcome; thus adding most detectors one to a new rule will most often lead to a rule set with lower prior probability without improving the likelihood, while adding one to an existing rule most often reduce both prior and likelihood.) Thus this approach would lead to slow mixing.

For improved efficiency, we instead implemented a two-step addition transformation. First, the algorithm chooses a position in $R$ (either in an existing rule, or a new rule) and considers the suite $\mathcal{S}$ of alternative rule sets which could be formed by adding any detector from the pool $\mathcal{R}$ to that position. A transition from $R$ into $\mathcal{S}$ is proposed, marginalizing over $\mathcal{S}$; then, if that transition is accepted, the algorithm chooses a particular $R'$ in $\mathcal{S}$ to move to. The detector removal transition corresponds to the reverse of this step.

A similar efficiency argument motivates the transition which allows the movement of detectors between rules, which evaluates all possible positions in the rule set $R$ to which a chosen detector could be moved, rather than proposing a single move and accepting or rejecting it.

These transition steps are described in detail in the following sections.

##### 1.2.1 Note on marginalization of the regression coefficients

Rule sets $R$ are sampled with the regression coefficients $\boldsymbol{\beta}$ marginalized out:

$$\Pr(\mathbf{y}|\boldsymbol{\omega}, A) = \Pr(\mathbf{z}|\boldsymbol{\omega}, A) \propto \exp\left\{-\frac{1}{2}\mathbf{z}^T\left(\Omega^{-1} + \sigma_\beta^2 A(R)A(R)^T\right)\mathbf{z}\right\}. \quad (1.3)$$

The conditional posterior distribution of $R$ is then

$$\pi(R|\boldsymbol{\omega}, \mathbf{y}) \propto \pi(R)\exp\left\{-\frac{1}{2}\mathbf{z}^T\left(\Omega^{-1} + \sigma_\beta^2 A(R)A(R)^T\right)\mathbf{z}\right\}. \quad (1.4)$$

When a new $R$ is chosen, a new value for $\boldsymbol{\beta}$ is drawn from the new conditional distribution of $\boldsymbol{\beta}$ given $A(R)$ and $\boldsymbol{\omega}$.

##### 1.2.2 Additional notation

Let $r^{(k)}$ be the $k$-th detector in $\mathcal{D}$ (in the order established above), as distinct from $r_{ij}$, the $j$-th detector in rule $i$ within some rule set $R$ (where the detectors have been sorted within each rule according to that same ordering.) In some contexts we will define a rule set $R'$. Let $\rho'_i$ be the $i$th rule within $R'$, and $r'_{ij}$ be the $j$-th detector in $\rho'_i$, and so on for $\rho^*_i$, $r^*_{ij}$ and $R^*$ where appropriate, etc. Similarly let $z'^{(i)}$ be the representation of rule $i$ within rule set $R'$ as a vector.

Let $S$ be the operator which sorts a tuple of detectors according to the chosen ordering on $\mathcal{D}$.

##### 1.2.3 Detector replacement

In this step, we propose the replacement of a detector in the rule set with some other detector in $\mathcal{D}$. For efficiency, we consider at once all possible options: the existing rule set $R$, and the $n_\mathcal{D} - 1$ possible rule sets that may be formed by replacing a particular detector in $R$ with an alternative.

In detail, we choose a detector in $r_{ij}$ in $R$ at random (with equal probability over the collection of all detectors in all rules) and obtain a new rule list $R'$ by deleting $r_{ij}$ from $R$, that is,

$$R' = \big(\rho_1, \ldots, (r_{i1}, \ldots, r_{i(j-1)}, r_{i(j+1)}, \ldots, r_{in_i}), \ldots, \rho_m\big) \,.$$

Then we form a set of of rule sets

$$\begin{aligned}
\Delta &= \Big\{R'_1, \ldots, R'_{n_\mathcal{D}}\Big\} \\
&= \Big\{ \Big(\rho_1, \ldots, S(r'_{i1}, \ldots, r'_{in'_i}, r^{(1)}), \ldots, \rho_m\Big) \,, \\
&\qquad \Big(\rho_1, \ldots, S(r'_{i1}, \ldots, r'_{in'_i}, r^{(2)}), \ldots, \rho_m\Big) \,, \\
&\qquad\qquad \vdots \\
&\qquad \Big(\rho_1, \ldots, S(r'_{i1}, \ldots, r'_{in'_i}, r^{(n_\mathcal{D})}), \ldots, \rho_m\Big) \Big\}
\end{aligned}$$

that is, the set of rule sets resulting from every possible replacement of $r_{ij}$ with any element of $\mathcal{D}$. Note that while $R'$ may not be a valid rule set (because $\rho'_i$ may be empty) each element of $\Delta$ is valid, and furthermore $R \in \Delta$.

Finally, we choose a new state from among $\{R'_1, \ldots, R'_{n_\mathcal{D}}\}$ with probabilities proportional to

$$(z'^{(i)}_1 + 1)\pi(R'_1|\boldsymbol{\omega}, \mathbf{y}), (z'^{(i)}_2 + 1)\pi(R'_2|\boldsymbol{\omega}, \mathbf{y}), \ldots, (z'^{(i)}_{n_\mathcal{D}} + 1)\pi(R'_{n_\mathcal{D}}|\boldsymbol{\omega}, \mathbf{y})$$

(see section 2.1, below.)

##### 1.2.4 Addition of a detector

For improved efficiency, we split our consideration of the addition proposal into two steps. Having chosen a position in the rule set at which to add a detector, we first decide whether or not to add a detector at all; if that proposal is accepted, we then choose from among the possible detectors to add.

The details of the procedure depend on whether we are adding a detector by creating a new length-1 rule in $R$, or adding a detector to an existing rule in $R$. We choose the position where the detector will be added from the following options:

- In a new, length-1 rule before the first rule in $R$, or after an existing rule in $R$ ($m + 1$ options)
- Into an existing rule in $R$ ($m$ options).

We consider each possible position equally likely, so that we propose insertion in a new rule with probability $(m+1)/(2m+1)$ and insertion into an existing rule with probability $m/(2m + 1)$.

**Addition of a new rule** Suppose that we have chosen to propose inserting the new rule at the $i$th position in the rule set. We define the set of possible rule sets we may form in this way as:

$$\begin{aligned}\mathcal{S}^* &= \left\{R'_1, \ldots, R'_{n_{\mathcal{D}}}\right\} \\ &= \Big\{ \left(\rho_1, \ldots, \rho_{i-1}, (r^{(1)}), \rho_i, \ldots \rho_m\right), \\ &\qquad\qquad \vdots \\ &\qquad \left(\rho_1, \ldots, \rho_{i-1}, (r^{(n_{\mathcal{D}})}), \rho_i, \ldots \rho_m\right) \Big\}\end{aligned}$$

As derived below (section 2.2.1), we accept the transition into $\mathcal{S}^*$ with probability

$$\max\left(1, \frac{2m+1}{1+\sum_{i=1}^{m} n_i} \frac{\pi_{S'}}{\pi_S(R)} \frac{L(\mathcal{S}^*|\boldsymbol{\omega})}{L(R|\boldsymbol{\omega})}\right),$$

where $\pi_{S'}$ is the structure contribution to the rule set prior $\pi_S(R'_k)$ shared by all $R'_k \in \mathcal{S}^*$.

If the proposal is accepted, we choose a particular $R'$ from $\mathcal{S}^*$ according to the conditional distribution over rule lists (1.4) restricted to $\mathcal{S}^*$.

**Addition to an existing rule** If instead we have chosen to propose adding a detector to existing rule $i^*$, the set of possible new rule sets we may form in this way is instead

$$\begin{aligned}\mathcal{S}^* &= \left\{R'_1, \ldots, R'_{n_\mathcal{D}}\right\} \\ &= \left\{ \left(\rho_1, \ldots, S(r_{i^*1}, \ldots, r_{i^*n_{i^*}}, r^{(1)}), \ldots \rho_m\right), \right. \\ &\qquad \vdots \\ &\qquad \left. \left(\rho_1, \ldots, S(r_{i^*1}, \ldots, r_{i^*n_{i^*}}, r^{(n_\mathcal{D})}), \ldots \rho_m\right) \right\}\end{aligned}$$

We ultimately accept the transition into $\mathcal{S}^*$ with probability

$$\max\left(1, T(\mathcal{S}^*)(2m+1)C_2 \frac{L(\mathcal{S}^*|\boldsymbol{\omega})}{L(R|\boldsymbol{\omega})}\right);$$

this probability is derived, and $T(\mathcal{S}^*)$ and $C_2$ defined, in section 2.2.2 below. If the move to $\mathcal{S}^*$ is accepted, we choose a particular $R'$ from $\mathcal{S}^*$ according to the conditional distribution over rule sets (1.4) restricted to $\mathcal{S}^*$.

##### 1.2.5 Removal of a detector

We treat each removal step as the reverse of one of the addition steps described above: that is, we consider it as a transition back to a specific shorter rule set from the space of possible rule sets obtained by addition of a detector to a particular position in that shorter rule set, without regard to which particular rule set in that space we happen to be visiting at the moment we consider the removal.

The details of the procedure depend on whether we are removing a detector from a length-1 rule in $R$ (and the length-1 rule itself), or removing a detector from a rule of length greater than 1. We choose a detector $r_{ij}$ at random from all the detectors in all the rules in $R$. Let

$$R^- = \begin{cases} \left(\rho_1, \ldots, \rho_{i-1}, \rho_{i+1}, \ldots, \rho_m\right), & n_i = 1 \quad \text{(case 1)} \\ \left(\rho_1, \ldots, (r_{i1}, \ldots, r_{i(j-1)}, r_{i(j+1)}, \ldots, r_{in_i}), \ldots, \rho_m\right), & n_i > 1 \quad \text{(case 2)} \end{cases}$$

We treat the removal step as the reverse of the addition step necessary to obtain $R$ from $R^-$, in that we consider it as a proposed transition back to

$R^-$ from the set $\mathcal{S}^* \ni R$ formed by adding each detector in $\mathcal{D}$ to the relevant position in $R^-$. Specifically, in case 1:

$$\begin{aligned}\mathcal{S}^* &= \left\{R'_1, \ldots, R'_{n_\mathcal{D}}\right\} \\ &= \Big\{ \left(\rho_1, \ldots, \rho_{i-1}, (r^{(1)}), \rho_i, \ldots \rho_m\right), \\ &\qquad\qquad \vdots \\ &\qquad \left(\rho_1, \ldots, \rho_{i-1}, (r^{(n_\mathcal{D})}), \rho_i, \ldots \rho_m\right) \Big\}\end{aligned}$$

A derivation exactly analogous to the treatment of the corresponding addition step shows that we should accept the proposed move from $\mathcal{S}^*$ to $R^-$ with probability

$$\max\left(1, \frac{\sum_{i=1}^m n_i}{2m-1} \frac{1}{C'_1} \frac{L(R^-|\boldsymbol{\omega})}{L(\mathcal{S}^*|\boldsymbol{\omega})}\right)$$

where

$$C'_1 = \frac{\pi_{S'}}{\pi_S(R^{-1})}$$

and $\pi_{S'}$ is the structure contribution to the rule set prior for any element of $\mathcal{S}^*$.

In case 2, for consistency with the addition step, denote the detector chosen for removal as $r_{i^*j}$ instead, and suppose $r_{i^*j} = r^{(k)}$. Then

$$\begin{aligned}\mathcal{S}^* &= \left\{R'_1, \ldots, R'_{n_\mathcal{D}}\right\} \\ &= \Big\{ \left(\rho_1, \ldots, S(r_{i^*1}, \ldots, r_{i^*n_{i^*}}, r^{(1)}), \ldots \rho_m\right), \\ &\qquad\qquad \vdots \\ &\qquad \left(\rho_1, \ldots, S(r_{i^*1}, \ldots, r_{i^*n_{i^*}}, r^{(n_\mathcal{D})}), \ldots \rho_m\right) \Big\}\end{aligned}$$

and we proceed analogously to our treatment of the corresponding addition step (see section 2.3). We accept the proposed move from $\mathcal{S}^*$ to $R^-$ with probability

$$\max\left(1, \frac{1}{2m+1} \frac{\sum_i n_i}{z_k^{(i)}} \frac{1}{C'_2} \frac{L(R^-|\boldsymbol{\omega})}{L(\mathcal{S}^*|\boldsymbol{\omega})}\right),$$

where

$$C'_2 = \frac{P_n(n_{i^*}-1)}{P_n(n_{i^*}-2)} \sum_{k=1}^{n_\mathcal{D}} \frac{M'^{(k)}_{i^*}}{M^-_{i^*}} p(r^{(k)}),$$

and $M^-_{i^*}$ is the multinomial coefficient associated with rule $i^*$ in $R^-$.

##### 1.2.6 Detector moves

Consider the two rule sets $R_a = ((r_1), (r_2))$ and $R_b = ((r1, r2))$. We are interested in discriminating between nonlinear interactions, as in $R_b$, and additive effects, as in $R_a$, so it is particularly important that the sampler can explore both types of rule efficiently. However, in a case where both $R_a$ and $R_b$ are high-likelihood models, moving from $R_a$ to $R_b$ by addition and removal of detectors alone may require a relatively low-likelihood intermediate step: e.g. $R_1 = ((r_1))$ or $R_2 = ((r_2))$ (or a low prior probability intermediate step, e.g. $R_3 = ((r_1, r_2), (r_1))$.) Thus this transition will typically occur infrequently.

To improve mixing in these cases we introduce a step which moves a detector from one primitive rule to another, allowing transition directly from $R_a$ to $R_b$, for example.

In detail, if the rule list is empty or contains only one detector, we do nothing. Otherwise we choose a detector $r_{ij}$ in $R$ at random (with equal probability over the collection of all detectors in all rules) and consider moving $r_{ij}$ (or an inverted version of $r_{ij}$, replacing 'above' with 'below' or vice-versa) to every rule in $R$, or to a new length-1 rule at each possible position in $R$.

We define

$$R' = \begin{cases} (\rho_1, \ldots, \rho_{i-1}, \rho_{i+1}, \ldots, \rho_m), & n_i = 1 \\ \big(\rho_1, \ldots, (r_{i1}, \ldots, r_{i(j-1)}, r_{i(j+1)}, \ldots, r_{in_i}), \ldots, \rho_m\big), & n_i > 1 \end{cases}$$

and let $r^+ = r_{ij}$ and let $r^-$ be the detector identical to $r^+$ but with opposite direction (i.e., replacing 'above' with 'below' or vice-versa) which is guaranteed to also be included in the population $\mathcal{D}$. Define the following sets of rules:

$$\Delta_1^+ = \Big\{ \big((r^+), \rho'_1, \ldots, \rho'_{m'}\big), \\ \big(\rho'_1, (r^+), \rho'_2 \ldots, \rho'_{m'}\big), \\ \vdots \\ \big(\rho'_1, \ldots, \rho'_{m'}, (r^+)\big) \Big\}$$

$$\Delta_1^- = \Big\{ \big((r^-), \rho'_1, \ldots, \rho'_{m'}\big), \\ \big(\rho'_1, (r^-), \rho'_2 \ldots, \rho'_{m'}\big), \\ \vdots \\ \big(\rho'_1, \ldots, \rho'_{m'}, (r^-)\big) \Big\}$$

$$\Delta_2^+ = \Big\{ \Big(S(r'_{11}, \ldots, r'_{1n'_1}, r^+) \ldots, \rho'_{m'}\Big), \\ \vdots \\ \Big(\rho'_1, \ldots, S(r'_{m'1}, \ldots, r'_{m'n'_{m'}}, r^+)\Big) \Big\}$$

$$\Delta_2^- = \Big\{ \Big(S(r'_{11}, \ldots, r'_{1n'_1}, r^-) \ldots, \rho'_{m'}\Big), \\ \vdots \\ \Big(\rho'_1, \ldots, S(r'_{m'1}, \ldots, r'_{m'n'_{m'}}, r^-)\Big) \Big\}$$

Let $\Delta = \Delta_1^+ \cup \Delta_1^- \cup \Delta_2^+ \cup \Delta_2^-$. (Note $R \in \Delta$.) Then choose a new value for $R$ from the posterior distribution of $R$ conditional on $\omega$ restricted to the set $\Delta$, using (1.4) to evaluate the unnormalized conditional probabilities of the elements of $\Delta$.

#### 1.3 Sampling parameters of the prior distribution

Exploiting conjugacy, we draw new values for $\Theta_0$ from $\mathrm{Beta}(a_\Theta + 1, b_\Theta)$ if $m = 0$ (the rule list is empty) or $\mathrm{Beta}(a_\Theta, b_\Theta + 1)$ if $m \geq 1$ (the rule list is nonempty.)

The phylogenetic prior parameters $\sigma_L$ and $\mu_L$ are updated through simple Metropolis-Hastings updates. We propose $\sigma'_L = \|\sigma_L + \delta_\sigma\|$, $\mu'_L = \|\mu_L + \delta_\mu\|$, drawing $\delta_\mu$ from $N(0, \Delta_L^2/4)$ and $\delta_\sigma$ from a $N(0, \Delta_L/2)$.

Similarly, for the time window prior parameters $f_w$ and $c_w$, we propose $f'_w = \|((f_w + \delta_f + 1) \mod 2) - 1\|$ (so that increments which would cross outside of the interval $[0, 1]$ instead are reflected off its boundaries) and $c'_w = \|c_w + \delta_w\|$, drawing $\delta_f$ from a Gaussian of mean 0 and standard deviation 0.2 and $\delta_w$ from a Gaussian of mean 0 and standard deviation $0.2c_{w,\text{typical}}$.

#### 1.4 Initialization

For the initial state of each Markov chain we set $R$ to a rule set with one rule containing a single detector, chosen from the detector population to minimize the Hamming distance between the detector's vector of truth values and the true outcome vector (choosing arbitrarily in case of ties.) Then we set each element of $\boldsymbol{\omega}$ to 1.0, each element of $\boldsymbol{\beta}$ to 0.0, $\mu_L$ to $\Lambda_L$, and $\sigma_L$ to $5\Delta_L$ (unless the upper bound on the range of values allowed under the uniform prior on $\sigma_L$ has been set to less than this, in which case we set $\sigma_L$ to its upper bound.)

### Chapter 2

### Detailed derivation of MCMC acceptance probabilities

Here we derive in detail the acceptance probabilities for the MCMC steps outlined above.

#### 2.1 Detector replacement

Recall that in this step we must choose a new state from the set of of rule sets

$$\begin{aligned}\Delta &= \left\{R'_1, \ldots, R'_{n_{\mathcal{D}}}\right\} \\ &= \Big\{ \left(\rho_1, \ldots, S(r'_{i1}, \ldots, r'_{in'_i}, r^{(1)}), \ldots, \rho_m\right), \\ &\qquad \left(\rho_1, \ldots, S(r'_{i1}, \ldots, r'_{in'_i}, r^{(2)}), \ldots, \rho_m\right), \\ &\qquad\qquad \vdots \\ &\qquad \left(\rho_1, \ldots, S(r'_{i1}, \ldots, r'_{in'_i}, r^{(n_{\mathcal{D}})}), \ldots, \rho_m\right) \Big\}\end{aligned}$$

that is, the set of rule sets resulting from every possible replacement of $r_{ij}$ with any element of $\mathcal{D}$.

We would like to choose a new value for $R$ from the posterior distribution of $R$, conditional on $\omega$, restricted to the set $\Delta$. It is straightforward to evaluate the unnormalized conditional probabilities

$$\pi(R'_1|\boldsymbol{\omega}, \mathbf{y}), \pi(R'_2|\boldsymbol{\omega}, \mathbf{y}), \ldots, \pi(R'_{n_{\mathcal{R}}}|\boldsymbol{\omega}, \mathbf{y})$$

using (1.4).

However, if we simply choose from $\Delta$ according to those probabilities, we may violate detailed balance. Suppose, having chosen $R'_k$ as the new value of $R$, we immediately perform another detector update of this type. What is the probability that we consider again this *exact* transition, that is, choose again from the same set $\Delta$? This occurs if and only if the detector we choose to update is a copy of $r^{(k)}$ in rule $i$, which it will be with probability equal to the number $z_k^{(i)}$ of copies of detector $r^{(k)}$ in rule $i$, divided by the total number of detectors in $R$.

All elements of $\Delta$ contain the same total number of detectors, but they may differ in the final number of copies present in rule $i$ of the primitive we have added to $R'$ to obtain them: ordering the elements of $\Delta$ as above, these numbers will be $z_1'^{(i)} + 1, z_2'^{(i)} + 1, \ldots, z_{n_\mathcal{D}}'^{(i)} + 1$. By choosing the new state from among $\{R'_1, \ldots, R'_{n_\mathcal{D}}\}$ with probabilities proportional to

$$(z_1'^{(i)} + 1)\pi(R'_1|\boldsymbol{\omega}, \mathbf{y}), (z_2'^{(i)} + 1)\pi(R'_2|\boldsymbol{\omega}, \mathbf{y}), \ldots, (z_{n_\mathcal{D}}'^{(i)} + 1)\pi(R'_{n_\mathcal{D}}|\boldsymbol{\omega}, \mathbf{y})$$

we preserve detailed balance.

#### 2.2 Addition of detectors

##### 2.2.1 As new rules

Recall that, having chosen to propose inserting the new rule at the $i$th position in the rule set, we defined the set of possible rule sets we might form in this way as:

$$\begin{aligned}
\mathcal{S}^* &= \left\{ R'_1, \ldots, R'_{n_\mathcal{D}} \right\} \\
&= \Big\{ \left(\rho_1, \ldots, \rho_{i-1}, (r^{(1)}), \rho_i, \ldots \rho_m\right), \\
&\qquad\qquad \vdots \\
&\qquad \left(\rho_1, \ldots, \rho_{i-1}, (r^{(n_\mathcal{D})}), \rho_i, \ldots \rho_m\right) \Big\}
\end{aligned}$$

With what probability should we accept the transition into $\mathcal{S}^*$? Let $m' = m + 1$ be the common length of the rule sets in $\mathcal{S}$. The prior probability of a candidate rule set $R'_k$ in $\mathcal{S}^*$ is

$$\begin{aligned}
\pi(R'_k) &= (1-\Theta_0)\, P_m(m'-1) \left( \prod_{i=1}^{m} P_n(n_i - 1) M_i p(r_{i1}) p(r_{i2}) \ldots p(r_{in_i}) \right) P_n(0) p(r^{(k)}) \\
&= \pi_S(R'_k) \left( \prod_{i=1}^{m} M_i p(r_{i1}) p(r_{i2}) \ldots p(r_{in_i}) \right) p(r^{(k)}) \\
&= \frac{\pi_S(R'_k)}{\pi_S(R)} \pi_S(R) \left( \prod_{i=1}^{m} M_i p(r_{i1}) p(r_{i2}) \ldots p(r_{in_i}) \right) p(r^{(k)}) \\
&= \frac{\pi_S(R'_k)}{\pi_S(R)} \pi(R) p(r^{(k)})
\end{aligned}$$

The structure contribution to the rule set prior $\pi_S(R'_k)$ is the same for all $R'_k \in \mathcal{S}^*$; denote its value by $\pi_{S'}$.

It follows

$$\begin{aligned}
\frac{\pi(\mathcal{S}^*)}{\pi(R)} &= \sum_{R' \in \mathcal{S}^*} \frac{\pi(R')}{\pi(R)} \\
&= \frac{\pi_{S'}}{\pi_S(R)} \sum_{k=1}^{n_{\mathcal{D}}} p(r^{(k)}) \\
&= \frac{\pi_{S'}}{\pi_S(R)} \\
&\equiv C_1.
\end{aligned}$$

We can evaluate the likelihood of $\mathbf{y}$ given $R$ and $\boldsymbol{\omega}$ using (1.3); call this $L(R|\boldsymbol{\omega})$. The likelihood of $\mathbf{y}$ given $\omega$ and the condition that the rule set is some element of $\mathcal{S}^*$ is

$$L(\mathcal{S}^*|\boldsymbol{\omega}) = \sum_{R' \in \mathcal{S}^*} \frac{\pi(R')}{\pi(\mathcal{S}^*)} L(R'|\boldsymbol{\omega}) = \sum_{k=1}^{n_{\mathcal{D}}} p(r^{(k)}) L(R'_k|\boldsymbol{\omega})$$

and each term in the sum we can likewise evaluate using (1.3). We accept the transition into $\mathcal{S}^*$ with probability

$$\max\left(1, \frac{2m+1}{1+\sum_{i=1}^{m} n_i} C_1 \frac{L(\mathcal{S}^*|\boldsymbol{\omega})}{L(R|\boldsymbol{\omega})}\right).$$

The term

$$\frac{2m+1}{1+\sum_{i=1}^{m} n_i},$$

whose denominator is the total number of detectors in the proposed new rule list, maintains detailed balance by accounting for the difference between the probability of choosing the particular position at which we have proposed to add a detector to $R$ and the probability of proposing to remove the detector we have just added from the new rule set $R'$.

Note that, if the proposal is accepted, it is straightforward to choose a particular $R'$ from $\mathcal{S}^*$ according to the conditional distribution over rule sets (1.4) restricted to $\mathcal{S}^*$ because we have already calculated the (nonnormalized) probabilities of each option,

$$p(r^{(k)})L(R'_k|\boldsymbol{\omega})$$

above.

##### 2.2.2 To existing rules

Recall that, having chosen to propose adding a detector to existing rule $i^*$, we defined the set of possible new rule sets we might form in that way as

$$\begin{aligned}
\mathcal{S}^* &= \left\{R'_1, \dots, R'_{n_\mathcal{D}}\right\} \\
&= \Big\{ \left(\rho_1, \dots, S(r_{i^*1}, \dots, r_{i^*n_{i^*}}, r^{(1)}), \dots \rho_m\right), \\
&\qquad\qquad\qquad \vdots \\
&\qquad \left(\rho_1, \dots, S(r_{i^*1}, \dots, r_{i^*n_{i^*}}, r^{(n_\mathcal{D})}), \dots \rho_m\right) \Big\}
\end{aligned}$$

With what probability should we accept the transition into $\mathcal{S}^*$? Here,

$$\begin{aligned}
\pi(R'_k) &= (1-\Theta_0)\, P_m(m-1) \left( \prod_{i \neq i^*} P_n(n_i - 1) M_i p(r_{i1}) p(r_{i2}) \dots p(r_{in_i}) \right) \cdot \\
&\qquad \left(P_n(n'_{i^*} - 1) M'_{i^*} p(r_{i^*1}) p(r_{i^*2}) \dots p(r_{i^*n_{i^*}}) p(r^{(k)})\right) \\
&= \pi(R) \frac{P_n(n'_{i^*} - 1)}{P_n(n_{i^*} - 1)} \frac{M'_{i^*}}{M_{i^*}} p(r^{(k)}).
\end{aligned}$$

Let $M'^{(k)}_{i^*}$ be the multinomial coefficient associated with rule $i^*$ in $R'_k \in \mathcal{S}^*$. Then

$$\begin{aligned}\frac{\pi(\mathcal{S}^*)}{\pi(R)} &= \sum_{R' \in \mathcal{S}^*} \frac{\pi(R')}{\pi(R)} \\ &= \frac{P_n(n'_{i^*} - 1)}{P_n(n_{i^*} - 1)} \sum_{k=1}^{n_{\mathcal{D}}} \frac{M'^{(k)}_{i^*}}{M_{i^*}} p(r^{(k)}) \\ &\equiv C_2\end{aligned}$$

Though less simple in form than $C_1$ this is straightforward to evaluate. Similarly, we may evaluate

$$L(\mathcal{S}^*|\boldsymbol{\omega}) = \sum_{R' \in \mathcal{S}^*} \frac{\pi(R')}{\pi(\mathcal{S}^*)} L(R'|\boldsymbol{\omega}) = \frac{1}{C_2} \frac{P_n(n'_{i^*} - 1)}{P_n(n_{i^*} - 1)} \sum_{k=1}^{n_{\mathcal{D}}} \frac{M'^{(k)}_{i^*}}{M_{i^*}} p(r^{(k)}) L(R'_k|\boldsymbol{\omega}).$$

In calculating the relative probabilities of proposing to add a detector to this particular rule and proposing the corresponding reverse move from $\mathcal{S}^*$ to $R$, we must note that if we add a detector $r^{(k)}$ to rule $i^*$ which already contains $z_k^{(i^*)}$ copies of $r^{(k)}$, the probability of proposing the reverse move is

$$\frac{z_k^{(i^*)} + 1}{1 + \sum_{i=1}^m n_i},$$

which is not constant across $\mathcal{S}^*$. Noting that

$$\begin{aligned}\frac{M'^{(k)}_{i^*}}{M_{i^*}} &= \frac{n_{i^*} + 1}{z_k^{(i^*)} + 1} \\ z_k^{(i^*)} + 1 &= (n_{i^*} + 1) \frac{M_{i^*}}{M'^{(k)}_{i^*}},\end{aligned}$$

we evaluate the acceptance probability using the *expected* probability $T$ of proposing the reverse move, under the posterior distribution restricted to $\mathcal{S}^*$.

$$\begin{aligned} T(\mathcal{S}^*) &= \frac{1}{1+\sum_{i=1}^m n_i}\left(\frac{\sum_{R'\in\mathcal{S}^*}\left(z_k^{(i^*)}+1\right)\pi(R')L(R'|\boldsymbol{\omega})}{\sum_{R'\in\mathcal{S}^*}\pi(R')L(R'|\boldsymbol{\omega})}\right) \\ &= \frac{(n_{i^*}+1)^{-1}}{1+\sum_{i=1}^m n_i}\left(\frac{\sum_{R'\in\mathcal{S}^*}(M_{i^*}/M_{i^*}'^{(k)})\pi(R')L(R'|\boldsymbol{\omega})}{\sum_{R'\in\mathcal{S}^*}\pi(R')L(R'|\boldsymbol{\omega})}\right) \\ &= \frac{(n_{i^*}+1)^{-1}}{1+\sum_{i=1}^m n_i}\left(\frac{\sum_{R'\in\mathcal{S}^*}p(r^{(k)})L(R'|\boldsymbol{\omega})}{\sum_{R'\in\mathcal{S}^*}(M_{i^*}'^{(k)}/M_{i^*})p(r^{(k)})L(R'|\boldsymbol{\omega})}\right) \end{aligned}$$

and ultimately accept the transition into $\mathcal{S}^*$ with probability

$$\max\left(1, T(\mathcal{S}^*)(2m+1)C_2\frac{L(\mathcal{S}^*|\boldsymbol{\omega})}{L(R|\boldsymbol{\omega})}\right),$$

Note that, as in the previous case, if the move to $\mathcal{S}^*$ is accepted, it is straightforward to choose a particular $R'$ from $\mathcal{S}^*$ according to the conditional distribution over rule sets (1.4) restricted to $\mathcal{S}^*$: here, the (nonnormalized) probabilities of each option are

$$\frac{M_{i^*}'^{(k)}}{M_{i^*}}p(r^{(k)})L(R_k'|\boldsymbol{\omega}).$$

#### 2.3 Removal of detectors, case 2

Note that the probability of proposing this or an equivalent removal step (given that we will propose to remove a detector from $R$) is

$$\frac{z_k^{(i^*)}}{\sum_i n_i}$$

(here we are summing the lengths of the rules in $R$, and counting the number of copies of detector $k$ in the $i^*$-th rule of $R$, *not* in $R^-$.) The probability of proposing a transition from $R^-$ to $\mathcal{S}^*$ (given that we will propose to add a detector to $R^-$) is $1/(2m+1)$.

### Chapter 3

### Parameter sensitivity assessment

Chapter 1 of the Supplementary Note provides default values for the various parameters and hyperparameters required to specify the prior distribution over rule sets. To evaluate the sensitivity of the performance of the MITRE algorithm to the choice of these parameters, we adjusted individual parameters or small groups of related parameters away from their default settings and applied MITRE to predict membership in the Russian cohort of the Vatanen et al. (2015) study. This dataset and prediction task were chosen to ensure robust cross-validation performance assessment when evaluating parameter sensitivity, because this case provided the largest number of subjects out of the available datasets. Performance was assessed using the F1 score calculated on held-out data across five folds of crossvalidation.

Hyperparameter values were assessed over a range of settings, including fairly extreme values as described below:

**Rule set structure parameters** We considered 'strict' and 'relaxed' settings for for the negative binomial priors on the rule set structure. The 'strict' settings were

$$\begin{aligned}\alpha_m &= 0.4\\ \beta_m &= 8.0\\ \alpha_n &= 0.2\\ \beta_n &= 4.0\end{aligned}$$

leading to a mean number of rules 1.05 and mean number of detectors per rule 1.1. The 'relaxed' settings were

$$\alpha_m = 3.0$$
$$\beta_m = 2.0$$
$$\alpha_n = 2.0$$
$$\beta_n = 1.0$$

leading to a mean number of rules 2.5 and mean number of detectors per rule 3.0. (All expectations are given in the limit of high $m_{\max}$ and $n_{\max}$; in practice, for all calculations, we set $m_{\max} = n_{\max} = 10$.)

The 'strict' parameter set thus reduces the model's flexibility considerably, while the 'relaxed' parameter set might be expected to lead to a tendency to overfit.

**Time window hyperparameter** We varied $\lambda_w$ to settings corresponding to expected values of $c_w$ of 0.5 and 50. Recall that $c_w$ scales the parameters of a beta distribution; thus, a small window concentration parameter $c_w < 1$ will lead to a distribution peaked near 0 and 1, favoring very short or very long windows and making it difficult to detect dynamics on a medium time scale, while a large $c_w \gg 1$ will disfavor, e.g., conjunctions of rules of different lengths, by focusing on rules which apply to around a fraction of the overall experimental time window which is very close to the current typical value $f_w$.

**Phylogeny-related hyperparameters** We varied the parameters for the phylogeny prior to encompass a large range. Reall that these parameters have a natural scale set by the $\log L_i$ values (and $\Delta_L$, the range from the 2.5th to 97.5th percentiles of those values.) For the data set analyzed median($\log L_i$) $= -2.05$, so that perturbing $\Lambda_L$ by 1 or 2 represents a large change. Similarly, a variance of $500\Delta_L^2$ implies that $\mu_L$ will often be substantially larger or smaller than the $\log L_i$ values, etc.

The table below presents the results of our sensitivity analysis, demonstrating that MITRE predictions retain high recall and precision across a broad range of hyperparameter values. A fair amount of variability is seen across the folds of crossvalidation, which is not surprising as this dataset, though the largest of those studied here, is still relatively small. Considering this variation, we did not see any significant sensitivity to the broad hyperparameter settings chosen.

| parameter | default | adjusted value | F1 score (point) | F1 score (ensemble) |
|---|---|---|---|---|
| default values all parameters | N/A | N/A | 0.833 (0.769-0.8333) | 0.833 (0.667-909) |
| rule set structure prior | see Supplementary Note | ‘strict’ | 0.727 (0.727-0.857) | 0.909 (0.727-0.909) |
| | | ‘relaxed’ | 0.800 (0.769-0.833) | 0.909 (0.800-0.909) |
| coefficient variance $\sigma_b^2$ | 100 | 10 | 0.833 (0.667-0.857) | 0.909 (0.800-0.909) |
| | | 200 | 0.727 (0.667-0.800) | 0.800 (0.727-0.923) |
| time window parameter $\lambda_w$ | 0.2 | 0.02 | 0.769 (0.769-0.800) | 0.833 (0.800-0.833) |
| | | 2.0 | 0.769 (0.615-0.769) | 0.800 (0.769-0.833) |
| $\Lambda_L$ | $\text{median}(\log L_i)$ | $\text{median}(\log L_i) - 1$ | 0.727 (0.667-0.833) | 0.800 (0.727-0.909) |
| | | $\text{median}(\log L_i) + 2$ | 0.727 (0.667-0.833) | 0.800 (0.667-0.833) |
| $\mu_L$ variance | $50\Delta_L^2$ | $5\Delta_L^2$ | 0.800 (0.769-0.909) | 0.800 (0.800-0.800) |
| | | $500\Delta_L^2$ | 0.667 (0.667-0.727) | 0.769 (0.727-0.833) |
| $\sigma_L$ upper bound | $25\Delta_L$ | $2.5\Delta_L$ | 0.769 (0.727-0.800) | 0.909 (0.833-0.909) |
| | | $250\Delta_L^2$ | 0.769 (0.615-0.833) | 0.800 (0.800-0.909) |

Table 3.1: MITRE predictive performance with adjusted hyperparameters, as applied to the problem of predicting membership in the Russian cohort in Vatanen et al. (2015). F1 scores shown are the median value and interquartile range for the held-out subjects across five folds of crossvalidation. See text for descriptions of the ‘strict’ and ‘relaxed’ settings for the negative binomial priors governing the rule set structure. For definitions of other parameters, see Chapter 1 of the Supplementary Note.
